# Biasing HER4 Tyrosine Kinase Signaling with Antibodies: Induction of Cell Death by Antibody-Dependent HER4 Intracellular Domain Trafficking

**DOI:** 10.1101/2019.12.20.883819

**Authors:** Romain Lanotte, Véronique Garambois, Nadège Gaborit, Christel Larbouret, Astrid Musnier, Pierre Martineau, André Pèlegrin, Thierry Chardès

**Author notes:** Corresponding author Dr T. Chardès, Institut de Recherche en Cancérologie de Montpellier, Montpellier, 34298, France. **Conflicts of interest:** R. Lanotte, P. Martineau, A. Pèlegrin and T. Chardès are inventors of the pending patent “Antibodies having specificity to HER4 and uses thereof”, The other authors declare no conflict of interest.

## Abstract

HER4 isoforms have oncogenic or tumor suppressor functions depending on their susceptibility to proteolytic cleavage and HER4 Intracellular Domain (4ICD) translocation. Here, we report that the NRG1 tumor suppressor mechanism through the HER4 JMa/CYT1 isoform can be mimicked by the agonist anti-HER4 antibody C6. NRG1 induced cleavage of poly(ADP-ribose) polymerase (PARP) and sub-G1 DNA fragmentation, and also reduced the metabolic activity of HER3-negative/HER4-positive cervical (C-33A) and ovarian (COV318) cancer cells. This effect was confirmed in HER4 JMa/CYT1-, but not JMa/CYT2-transfected BT549 triple-negative breast cancer cells. NRG1 favored 4ICD cleavage and retention in mitochondria in JMa/CYT1-transfected BT549 cells, leading to Reactive Oxygen Species (ROS) production through mitochondrial depolarization. Similarly, the anti-HER4 antibody C6, which binds to a conformational epitope located on aa 575-592 and 605-620 of HER4 domain IV, induced 4ICD cleavage and retention in mitochondria, and mimicked NRG1-mediated effects on PARP cleavage, ROS production, and mitochondrial membrane depolarization in cancer cells. *In vivo*, C6 reduced growth of COV434 and HCC187 tumor cell xenografts in nude mice. Biasing 4ICD trafficking to mitochondria with anti-HER4 antibodies to mimic NRG1 suppressor functions could be an alternative anti-cancer strategy.

## 1. Introduction

The Human Epidermal growth factor Receptor’s family (HER or ErbB) includes four Receptor Tyrosine Kinases (RTK; EGFR/HER1, HER2, HER3 and HER4) that play roles in development and cancer. EGFR and HER2 role in cancer progression led to the development of monoclonal antibodies (mAbs) against these receptors, such as cetuximab and trastuzumab [1]. More recently, HER3 also has been considered a key player in tumor signaling and resistance to cancer drugs, leading to the development of anti-HER3 mAbs [2]. Conversely, results with antagonist mAbs against HER4 have been disappointing, and currently no anti-HER4 mAb is used in the clinic.

HER4 is unusual among HER members. Its biology is more complex and its role in cancer is still controversial. HER4 is expressed in various cancers, such as blastoma, breast, lung, melanoma, pancreas, gastric, colorectal, ovarian and bladder cancer [3]. However, the prognostic significance of HER4 expression in cancer remains unclear, particularly in breast cancer where HER4 has been alternatively described as an oncogene [4] and a tumor suppressor [5]. These opposite effects are explained by the existence of four HER4 isoforms at the cell surface, each with its own downstream signaling pathway [6]. These isoforms (JMa/CYT1, JMa/CYT2, JMb/CYT1 and JMb/CYT2) differ in their ExtraCellular Domain (ECD) and IntraCellular Domain (ICD). Following activation, JMa isoforms are cleaved by a two-step process, catalyzed by TACE and then γ-secretase and called Regulated Intramembrane Proteolysis (RIP), to release the HER4 ECD and ICD (4ICD) [7]. 4ICD translocates to the nucleus where it acts on gene transcription to control multiple cellular pathways (differentiation, migration, proliferation) [8]. Conversely, JMb isoforms are not cleaved and act as classical RTKs. HER4 isoforms acquire the cytoplasmic domain CYT1 or CYT2 by alternative splicing [9]. CYT2 isoforms can only induce phosphorylation of MAPK pathway components, whereas the 16-amino acid extension present only in CYT1 isoforms allows the activation of the MAPK and also of the PI3K pathway [10].

Most studies describe HER4 isoforms and their main ligand NRG1β1 (NRG1 hereinafter) as oncogenes. JMa/CYT1 and JMa/CYT2 are widely co-expressed. Conversely, expression of JMb variants seems to be restricted to some tissues [6]. In cancer, JMa/CYT1 and JMa/CYT2 have been associated with poor prognosis, due to 4ICD translocation to the nucleus [11]. JMa/CYT1 has been implicated in tumor progression [12], and JMa/CYT2 is considered the most oncogenic isoform. Indeed, CYT2 is more stable than CYT1 in the cytosol [13], and its nuclear location is more robust, with better transcriptional activity [14]. Moreover, CYT2 can activate hyperplasia-related pathways, such as Wnt, β-catenin, and KITENIN [15], and JMa/CYT2 homodimers are constitutively phosphorylated to promote ligand-independent growth [16]. Both isoforms support cancer cell proliferation by modulating numerous signaling pathways [17].

However, in breast cancer, CYT1 isoforms have been associated also with inhibition of cancer cell proliferation [18]. In the cytosol of breast cancer cells, 4ICD directly induces apoptosis from mitochondria through its BH3-only domain [19], explaining the better survival of patients with high cytosolic 4ICD expression [20]. As HER4 plays a role in tissue homeostasis [21], which requires regulation of proliferation and cell death [22], HER4 and 4ICD might also play a tumor suppressor function that could be modulated by NRG1. Indeed, the *ERBB4* promoter is hypermethylated in cancer, and HER4 re-expression using demethylating agents induces apoptosis of breast cancer cells after NRG1 stimulation [23]. In breast cancer, NRG1 and HER4 induce cell cycle arrest by activating JNK through BRCA1 [24], and 4ICD might be a mitotic checkpoint [25], regulating cell cycle progression.

As NRG1 is considered a potential tumor suppressor gene [26] and the Y1056 residue in HER4-CYT1 variants is essential for tumor suppression [27], we hypothesized that the HER4 JMa/CYT1-NRG1 axis has a tumor suppressor function by localizing 4ICD in mitochondria where it can induce apoptosis through its BH3-only domain [19]. We also hypothesized that this NRG1-4ICD pathway could be activated by biased agonist or positive allosteric modulator mAbs, as described for G-Protein Coupled Receptor (GPCR) targeting [28]. By inducing specific conformational changes in the targeted receptor, these mAbs can selectively modulate specific signaling pathways [29]. Here, we demonstrated that NRG1 induces HER4 JMa/CYT1-expressing cancer cell death through 4ICD retention in mitochondria, and that the anti-HER4 antibody C6 mimics these NRG1-induced effects, leading to growth inhibition of ovarian and breast cancer xenografts.

## 2. Materials and Methods

### 2.1. Cell cultures

C-33A cervical cancer cells were cultured in EMEM medium (Thermo Fisher, Waltham, MA) supplemented with 10% fetal bovine serum (FBS) and antibiotics (Thermo Fisher). COV318 and COV434 ovarian cancer cells, and HEK293T human embryonic kidney cells were cultured in DMEM/F-12, GlutaMAX medium (Thermo Fisher) with 10% FBS and antibiotics. HEK293T cells were used for epitope mapping and full-length IgG production. BT549 and HCC1187 triple negative breast cancer (TNBC) cells were cultured in RPMI 1640, GlutaMAX medium with 10% FBS and antibiotics. All cell lines were routinely analyzed for absence of mycoplasma, using the MycoAlert Kit (Lonza, Switzerland). NIH3T3 mouse embryonic fibroblasts were stably transfected with pCMV_HER4 that encodes the JMa/CYT1 isoform, as previously described [2], and were used for whole cell panning by phage display. Transfected NIH3T3 cells were cultured in DMEM/F-12, GlutaMAX medium with 10% FBS, antibiotics and 10 μg/ml hygromycin B (Thermo Fisher). All cell lines were obtained from American Type Culture Collection (ATCC) (Rockville, MD), and were maintained at 37°C in a humidified atmosphere with 5% CO_2_.

### 2.2. Recombinant Proteins and Constructs

Recombinant Fc-tagged human and mouse HER4 ECD, and human NRG1 extracellular domain were purchased from R&D Systems (Minneapolis, MN). The HER4 JMa/CYT1 human sequence was amplified by PCR and subcloned in the pcDNA3.1 vector (Invitrogen, Carlsbad, CA) to create pCMV_HER4, used for stable transfection of NIH3T3 cells. pEZY3_JMa/CYT1 and pEZY3_JMa/CYT2 were used for transient transfection of BT549 cells. Both plasmids were generated using the Gateway technology. JMa/CYT1 was PCR-amplified from pCMV_HER4 and subcloned in the pDONR221 vector (Invitrogen) to generate pDONR221-JMa/CYT1, using the Gateway BP Clonase II Enzyme Mix (Invitrogen). pDONR221-JMa/CYT2 was purchased from Life Technologies (clone ID: IOH81996). Both pDONR221 constructs were subcloned in the pEZ3 final vector (Addgene, Watertown, MA) using the Gateway LR Clonase II Enzyme Mix (Invitrogen). All constructs were verified by sequencing. Constructs for epitope mapping (HER4_WT, HER4_P1m, HER4_P2m, HER4_P3m and HER4_P4m) were N-terminally Flag-tagged to monitor antigen expression in cells. All reagents and resources are listed in Supplementary Table S1.

### 2.3. Phage display selection and production of full-length IgGs

Two phage display selection experiments were performed in parallel: the first using NRG1-stimulated JMa/CYT1-transfected cells (whole cell panning by phage display) and the second using recombinant human HER4 ECD as targets. For the first one, JMa/CYT1-transfected NIH3T3 cells were stimulated with 50 ng/ml NRG1 at 37°C for 15min (to activate HER4) and then incubated with the HUSCI proprietary scFv library at 4°C for 4h. The HUSCI library was pre-incubated with non-transfected NIH3T3 cells at 4°C for 2h, to remove irrelevant binders. This synthetic library uses a single framework optimized for high-level expression. The side-chain diversity was restricted to five amino acids (Y,N,D,G,S) and introduced into the six Complementarity Determining Regions (CDRs) with variable VH-CDR3 lengths at the positions corresponding to the most contributing residues of the paratope [30]. After incubation, cells were washed twenty times with PBS and phages were eluted with trypsin, and treated with TPCK (Thermo Fisher) at room temperature for 10min. Eluted phages were amplified in TG1 bacteria (Agilent, Santa Clara, CA), rescued with the KM13 helper phage (New England Biolabs, Omaha, NE), and used for two other rounds of selection. Eluted phages from the third round were amplified in HB2151 bacteria (Agilent). Approximatively 400 clones were picked from Petri dishes and cultured in 96-well plates. Clone binding to NRG1-stimulated JMa/CYT1-transfected cells versus non-stimulated or non-transfected cells was screened by flow cytometry. Briefly, cells were seeded in 96-well plates (5×10^4^ cells/well) and incubated with scFv on ice for 3h. After washes, scFv binding was detected with the AF647-conjugated anti-c-Myc antibody (MBL International, Woburn, MA). The cell median fluorescence intensity was measured by flow cytometry using Cyan flow cytometer (Cytek Biosciences, Fremont, CA). In the second approach, binding to soluble recombinant human HER4 was assessed by ELISA. Briefly, Nunc MaxiSorp plates (Thermo Fisher) were coated with 1 μg/ml human HER4, and scFv clones (i.e., HUSCI proprietary scFv library) were added at room temperature for 2h. After washes, scFv binding was detected with a peroxidase-conjugated anti-Myc antibody (Santa Cruz Biotechnology, Santa Cruz, CA)) and TMB solution (Thermo Fisher). The reaction was stopped by adding 1M sulfuric acid before absorbance measurement at 450 nm (Multiskan; Thermo Fisher). From both selections, the VH and VL of positive scFv clones were sequenced and subcloned in a backbone human IgG1 vector. All candidates were sequence-verified and the IgG1 format was used for testing antibody binding to HER4, as previously below.

IgGs were produced in transiently transfected HEK293T cells, using Polyethylenimine Linear transfection reagent (PEI) (Polysciences, Warrington, PA), in serum-free DMEM. Briefly, cells were seeded in 150 mm Petri dishes and allowed to grow to 70% confluence. Cells were transfected with 15 μg of antibody-encoding plasmids and 240 μg of PEI, in 150mM NaCl solution for 5h. Medium was then replaced by serum-free DMEM and cells were incubated at 37°C in humidified atmosphere at 5% CO_2_. Supernatants from HEK293T cells were harvested 5 days after transfection. Cells and debris were removed by centrifugation, and supernatants were buffered with sodium phosphate. After filtration, IgGs in supernatants were purified through a HiTrap protein A column (GE Healthcare, Chicago, IL). Antibodies were eluted with 0.1M glycine pH 2.7, immediately buffered with 1M Tris solution pH 9, and then dialyzed and concentrated in PBS.

### 2.4. HER4 JMa/CYT1- and JMa/CYT2-encoding plasmid transfection in BT549 and C-33A cells

To ensure uniformity of transfection among experimental conditions and experiments, BT549 and C-33A cells were seeded in 150 mm Petri dishes and grown for 24h. Cells were transfected at 70% confluence with 15 μg of JMa/CYT1 or JMa/CYT2 plasmid and 240 μg of PEI diluted in 150mM NaCl solution for 5h. Medium (10% FBS) was then replaced by 1% FBS-medium. Next day, transfected cells were seeded in 6-well or 12-well plates (10% FBS-medium) and grown for 10h. Cells were then starved in 1% FBS-medium for 12h before NRG1 stimulation or antibody treatment.

### 2.5. Flow cytometry analysis

Cell surface receptor expression was analyzed in all cell lines with the following antibodies: cetuximab (EGFR) (Merck), trastuzumab (HER2) (Roche), SGP1 (HER3) (Santa Cruz Biotechnology), and H4.77.16 (HER4) (Thermo Fisher). Cetuximab and trastuzumab were purchased from the pharmacy of the ICM hospital. Exponentially growing cells were harvested and resuspended in FACS buffer (PBS/1% FBS). 3×10^5^ cells were incubated with 10 μg/ml of primary antibody on ice for 1h30. For competition experiments, cells were incubated with 15 μg/ml of the selected anti-HER4 mAbs and NRG1 at various concentrations (0.1 to 100 nM; 3 to 3000 ng/ml) on ice for 1h30. After washes, cells were incubated with FITC-conjugated goat anti-human IgG (Fc specific) (Sigma St-Louis, MI) or FITC-conjugated goat anti-mouse IgG (H+L) (Millipore). The cell median fluorescence intensity was measured by flow cytometry (FL1 channel) (Gallios apparatus, Beckman Coulter, Fullerton, CA).

### 2.6. HER4 binding by ELISA

Nunc MaxiSorp plates (Thermo Fisher) were coated with 250 ng/ml human or mouse recombinant HER4 overnight. After saturation at 37°C with PBS/2% BSA for 2h, antibodies were diluted in PBS/0.1% Tween and added for 2h at 37°C. After washes, antibody binding was detected by incubation with peroxidase-conjugated goat anti-human IgG (F(ab’)2 fragment specific) (Jackson ImmunoResearch, West Grove, PA) and TMB solution. The reaction was stopped by adding 1M sulfuric acid, and absorbance was measured at 450 nm.

### 2.7. RT-PCR analysis of HER4 isoform expression

RNA from 10^6^ cells was extracted using the RNeasy Mini Kit (Qiagen, Germantown, MD) according to the manufacturer’s instructions. 1 μg RNA was reverse transcribed using SuperScript III Reverse Transcriptase (Thermo Fisher), according to the manufacturer’s instructions. PCR was then performed to amplify the full-length HER4 isoforms, using the Dream Taq Green PCR Master Mix (Thermo Fisher), 10 μM primers and 2.5 μl cDNA, with the following conditions: 10 min initial denaturation at 95°C, 40 cycles (95°C for 60sec, 56.9°C for 90 secs, and 72°C for 90 secs), and 5 min at 72°C for the final extension. 15 μl from each PCR sample was used for agarose gel electrophoresis and visualized under UV using G:BOX (Pxi 4, Syngene).

### 2.8. Mitochondrial activity measurement

Mitochondrial activity of C-33A cells was measured with the CellTiter 96 Aqueous One Solution Cell Proliferation Assay (MTS) (Promega, Madison, WI). Cells were seeded in 96-well plates (5000 cells/well) in 10% FBS-medium and left at 37°C for 24h. Cells were then starved in 1% FBS-medium for 12h before stimulation with 30 ng/ml NRG1, with or without 100 μg/ml of mAbs in 1% FBS-medium, at 37°C for 5 days. Alternatively, cells were only incubated with 100 μg/ml of mAbs in 10% FBS-medium for 5 days. Mitochondrial activity was then assessed by adding 20 μl MTS directly to the cells. After 2h incubation at 37°C, supernatants were centrifuged to remove cell debris and transferred into a new 96-well plate for absorbance reading at 490 nm. Data were presented as arbitrary units (a.u.) or normalized as percentage of the value in untreated or NRG1-stimulated cells.

### 2.9. Adenylate kinase releasing assay

Adenylate kinase release from damaged cells was measured using the ToxiLight Bioassay Kit (Lonza), according to the manufacturer’s instructions. HCC1187 cells were seeded in 96-well plates (10000 cells/well) in 10% FBS-medium and left at 37°C for 24h. Cells were then starved for 12h and stimulated with 30 ng/ml NRG1 in 1% FBS-medium at 37°C for 72h. Plates were then left at room temperature for 5min before transferring 20 μl of supernatant to each well of white flat bottom 96-well plates (Greiner, Austria). After addition of 100 μl of Detection Reagent to each well and incubation at room temperature for 5min, bioluminescence was measured with a PHERAstar FS reader (BMG Labtech, Germany). Data were presented as arbitrary units (a.u.).

### 2.10. ROS production

ROS level was measured using the DCFDA/H2DCFDA Cellular Reactive Oxygen Species Detection Assay Kit (Abcam, UK). JMa/CYT1-, JMa/CYT2- and Mock-transfected BT549 cells were seeded (2×10^4^ cells/well) in black transparent flat bottom 96-well plate (Greiner), in 10% FBS-medium and grown at 37°C for 24h. All experiments were performed in medium without phenol red. After starvation in 1% FBS-medium for 12h, cells were stimulated with 30 ng/ml NRG1 or incubated with 20 μg/ml mAbs in 1% FBS-medium at 37°C for 24h. In the positive control, cells were incubated with 100 μM Tert-Butyl HydroPeroxyde (TBHP) for 4h to induce oxidative stress. At 45 min before the end of the experiment, 50 μM DCFDA was directly added to the wells, and plates were put back at 37°C. Results were immediately acquired with a PHERAstar FS reader (BMG Labtech) using the end-point mode (Ex/Em = 485/535 nm). Data were presented as Relative Luminescence Units (RLUs) or as fold-change compared with untreated cells.

### 2.11. Mitochondrial Membrane Potential

Mitochondrial Membrane Potential changes in JMa/CYT1-, JMa/CYT2- and Mock-transfected BT549 cells were evaluated using the DIOC6 intracellular probe (Abcam). Cells were seeded in 6-well plates (3×10^5^ cells/well) in 10% FBS-medium and grown at 37°C for 24h. Cells were then starved for 12h, and stimulated with 30 ng/ml NRG1 or incubated with 20 μg/ml mAbs in 1% FBS-medium at 37°C for 24h. After PBS washes, cells were trypsinized and cell pellets were incubated with 20nM DIOC6. As positive control of mitochondrial depolarization, untreated cells (only medium) were incubated with 100 μM 2-[2-(3-Chlorophenyl)hydrazinylyidene]propanedinitrile (CCCP) and DIOC6. After 20 min-incubation in the dark at 37°C, cells were centrifuged and pellets were re-suspended in hot PBS. Fluorescence intensity was immediately measured using a Gallios flow cytometer (Beckman Coulter).

### 2.12. Subcellular fractionation of BT549 transfected cells

Around 4×10^7^ BT549 cells were seeded in 150 mm Petri dishes for each condition and grown in 10% FBS-medium at 37°C for 24h. Then, cells were transiently transfected with 15 μg of plasmids encoding JMa/CYT1 or JMa/CYT2 and 240 μg of PEI in 150mM NaCl solution, in 10% FBS-medium for 5h. Afterwards, medium was replaced with 1% FBS-medium for 12h. Cells were then stimulated or not with 30 ng/ml NRG1 at 37°C for 24h, or incubated with 20 μg/ml anti-HER4 antibodies in 1% FBS-medium, with or without 30 ng/ml NRG1 at 37°C for 6h. Cells were then washed twice in cold PBS, transferred in 15 ml Falcon tubes and centrifuged. All the following steps were performed on ice. Cell pellets were diluted in cold 1 ml Mitochondrial Isolation Buffer (MIB) (200 mM sucrose, 10 mM TRIS/MOPS, 1 mM TRIS/EGTA and proteases inhibitors) pH 7.4. Cell suspensions were then transferred into Nalgene tubes (Thermo Fisher) and immediately passed through a T-18 IKA Ultra Turrax homogenizer (Staufen, Germany) at speed 2.5 for 10-15 secs, to break the plasma membrane. Cell membrane lysis was checked under a microscope, and a second run was performed to reach approximatively 90% of lysis. An aliquot of 50 μl was saved as Whole Cell Lysate (WCL). Lysates were then centrifuged three times with MIB buffer at 600g and 4°C for 10min, and supernatants (containing the mitochondrial and cytosolic fractions) were stored. The supernatants from the three centrifugation steps were pooled in 15 ml Falcon tubes and centrifuged at 600g at 4°C for 10 min, to eliminate nuclear contaminants, and then centrifuged twice at 7500g at 4°C for 10 min to isolate mitochondria. Cell pellets containing mitochondria were diluted in 50 μl lysis buffer (60 mM TRIS pH 6.8, 10% glycerol (v/v), 1% SDS (v/v)) (i.e., mitochondrial fraction). Supernatants were centrifuged at 600g at 4°C for 10 min to eliminate mitochondrial residual contaminants, and precipitated with 4 volumes of cold acetone at −20°C for at least 1h. After several washes and centrifugations (15000g at 4°C for 10 min), pellets were dried and dissolved in 50 μl water (i.e., cytosol fraction). Protein concentration in all fractions was evaluated using a Nanodrop 2000 (Fisher Scientific). Samples were diluted in 2X Laemmli buffer and boiled at 95°C for 10 min. 30 μg of protein lysate from each fraction was analyzed by western blotting using the anti-HER4 antibody E200 (Abcam) to determine HER4/4ICD subcellular localization. An anti-alpha-tubulin antibody (Cell Signaling Technology) was used as control for the cytosolic fraction, and antibodies against TIM23 (BD Biosciences,) TFAM (Cell Signaling Technology) and VDAC1 (Millipore) as controls for the mitochondrial fraction.

### 2.13. Western blot analysis

Scraped cell samples were boiled in 1X lysis buffer (60 mM TRIS pH 6.8, 10% glycerol (v/v), 1% SDS (v/v)) at 95°C for 10 min, and protein concentration was determined using a Nanodrop 2000 (Fisher Scientific). Bromophenol blue was added, and 20-50 μg of protein lysates were separated by SDS-PAGE and then transferred onto PVDF membranes (GE Healthcare). Membranes were saturated in Tris Buffer Saline/0.1%Tween (TBS-T) and 5% non-fat dry milk with gentle shaking at room temperature for 2h. Membranes were incubated with rabbit or mouse primary antibodies at 4°C with gentle shaking overnight. After washes in TBS-T, membranes were incubated with the appropriate peroxidase-conjugated anti-mouse (Jackson ImmunoResearch) or anti-rabbit (Sigma) antibodies at room temperature for 1h. After washes in TBS-T, protein expression levels were detected with Western Lightening Ultra (Perkin Elmer) or SuperSignal West Femto Maximum Sensitivity Substrate (Thermo Fisher) and the G:BOX system (Pxi 4, Syngene). Data were analyzed using the ImageJ software (NIH), and are representative of at least two independent experiments.

### 2.14. Epitope Mapping

#### Design of HER4 mutants

The D5 and C6 epitopes were predicted using MabTope [31]. Briefly, MabTope is a docking-based method that generates 5×10^8^ poses for each antibody-antigen complex and filters these poses in order to obtain the 30 best solutions (Supplementary Fig. S5A), using several scoring functions. The interface analysis in these 30 top-ranked solutions allows identifying residues that exhibit the highest probability of being implicated in the interaction (Supplementary Fig. S5B). The D5 and C6 epitopes were predicted to be located in the same HER4 regions (P1 to P4) (Supplementary Fig. S5B). The amino acids that were identified as highly probable epitope residues were mutated to alanine to create one mutated HER4 construct for each initially identified region. With the aim of not affecting HER4 3D structure, prolines and other amino-acid residues the lateral chains of which are not exposed at the HER4 surface were not mutated. The wild type (HER4_WT) and mutated constructs (HER4_P1m, HER4_P2m, HER4_P3m and HER4_P4m) (Fig. 4C) were N-terminally Flag-tagged to monitor antigen expression in cells. Gene optimization, synthesis and cloning in pcDNA3.1+ were performed by GenScript (Piscataway, NJ).

#### Flow cytometry analysis of wild type and mutated HER4-transfected HEK293 cells

HEK293 cells were seeded in 10 cm dishes in 10% FBS-medium and grown at 37°C for 24h. Cells were then transiently transfected with 5 μg of each construct using Metafectene (Biontex Lab.; München, Germany), according to the manufacturer’s instructions. After 24h, cells were trypsinized, filtered through a 70 μm strainer (Laboteca; McAllen, TX) and fixed/permeabilized using the Cytofix/Cytoperm Solution Kit (BD Biosciences), according to the manufacturer’s instructions. 2×10^5^ cells/tube were incubated with 5 μg of C6 or D5 (anti-HER4 mAbs), or control antibody in 100 μl of Perm/Wash buffer at room temperature for 1h. Cells were then washed three times in 2 ml PBS supplemented with 2mM EDTA and 1% FBS. Cells were then co-incubated with 1 μg of APC-conjugated anti-IgG antibody (Miltenyi Biotec; Bergisch Gladbach, Germany) and 0.5 μg of PE-conjugated anti-Flag antibody (Miltenyi Biotec) in 100 μl of Perm/Wash buffer at room temperature for 45 min. Cells were first washed with 2 ml PBS-EDTA with 1% FBS, and then with 2 ml PBS-EDTA before suspension in 150 μl PBS-EDTA. Expression data were collected with a MACSQuant Analyzer 10 (Miltenyi Biotec) and analyzed using FlowJo (data from 3 independent experiments). The number of APC+PE+ double-positive cells was recorded for each sample and normalized on the total number of PE+ cells in the sample.

### 2.15. Tumor xenograft studies

All procedures were performed in compliance with the French regulations and ethical guidelines for experimental animal studies in an accredited establishment (Agreement No. C34-172-27). All experiments followed the relevant regulatory standards. Female Hsd nude mice were obtained from Envigo (Huntingdon, UK). At 6 weeks of age, mice were anesthetized and subcutaneously grafted with 8×10^6^ COV434 ovarian, 1×10^7^ C-33A cervical, or 1×10^7^ HCC1187 TNBC cells diluted in Matrigel solution (Corning, NY) in a final volume of 150 μl. Tumors were measured with a caliper twice a week to monitor size before and after treatment start. Once tumor size reached around 150 mm^3^, mice were randomly distributed in groups of 10 mice and treatments started (intraperitoneal injections): irrelevant antibody (20 mg/kg), anti-HER4 D5 (20 mg/kg) and anti-HER4 C6 (20 mg/kg) twice per week for 4 weeks, or carboplatin (60 mg/kg) once per week for 4 weeks.

### 2.16. Statistical analysis

All statistical tests were performed using the Prism software v5.1 (GraphPad). Comparisons between groups were performed using the two-tailed unpaired Student’s *t*-test. ANOVA was used for Figures 4D and S5. For all experiments, * is p<0.1, ** is p<0.01 and *** is p<0.001.

## 3. Results

### 3.1. NRG1 induces cell death in HER4-expressing C-33A and COV318 cancer cells

We first studied the involvement of the NRG1/HER4 axis in tumor suppression in C-33A (cervical) and COV318 (ovary) cancer cell lines that strongly express HER4, but not HER3. These two cell lines also express EGFR and HER2 at different levels, as shown by flow cytometry (Supplementary Fig. S1). Stimulation of C-33A cells with NRG1 induced strong poly(ADP-ribose) polymerase (PARP) cleavage over time, suggesting that cell death is triggered after HER4 activation by NRG1 (Fig. 1A). NRG1 also increased DNA fragmentation, as indicated by the finding that around 60% of NRG1-stimulated C-33A cells were in the sub-G1 phase at 48h-72h after addition of NRG1 (Fig. 1B). NRG1 also significantly decreased the metabolic activity (MTS assay) of C-33A cells (Fig. 1C), suggesting that NRG1-induced cell death occurs through mitochondria. Similarly, NRG1 stimulation of COV318 cells induced PARP cleavage (Fig. 1D) already at 24h-post NRG1 stimulation, and cell membrane disruption (Fig. 1E). We observed this “broken” phenotype also in HCC1187 TNBC cells that express HER4 (Supplementary Fig. S2A), as confirmed by adenylate kinase release increase in the extracellular medium in NRG1-stimulated HCC1187 cells compared with untreated cells (Supplementary Fig. S2B). These experiments demonstrated that the NRG1-HER4 axis induces cell death in C-33A, COV318 and HCC1187 cancer cells.

**Fig. 1.**
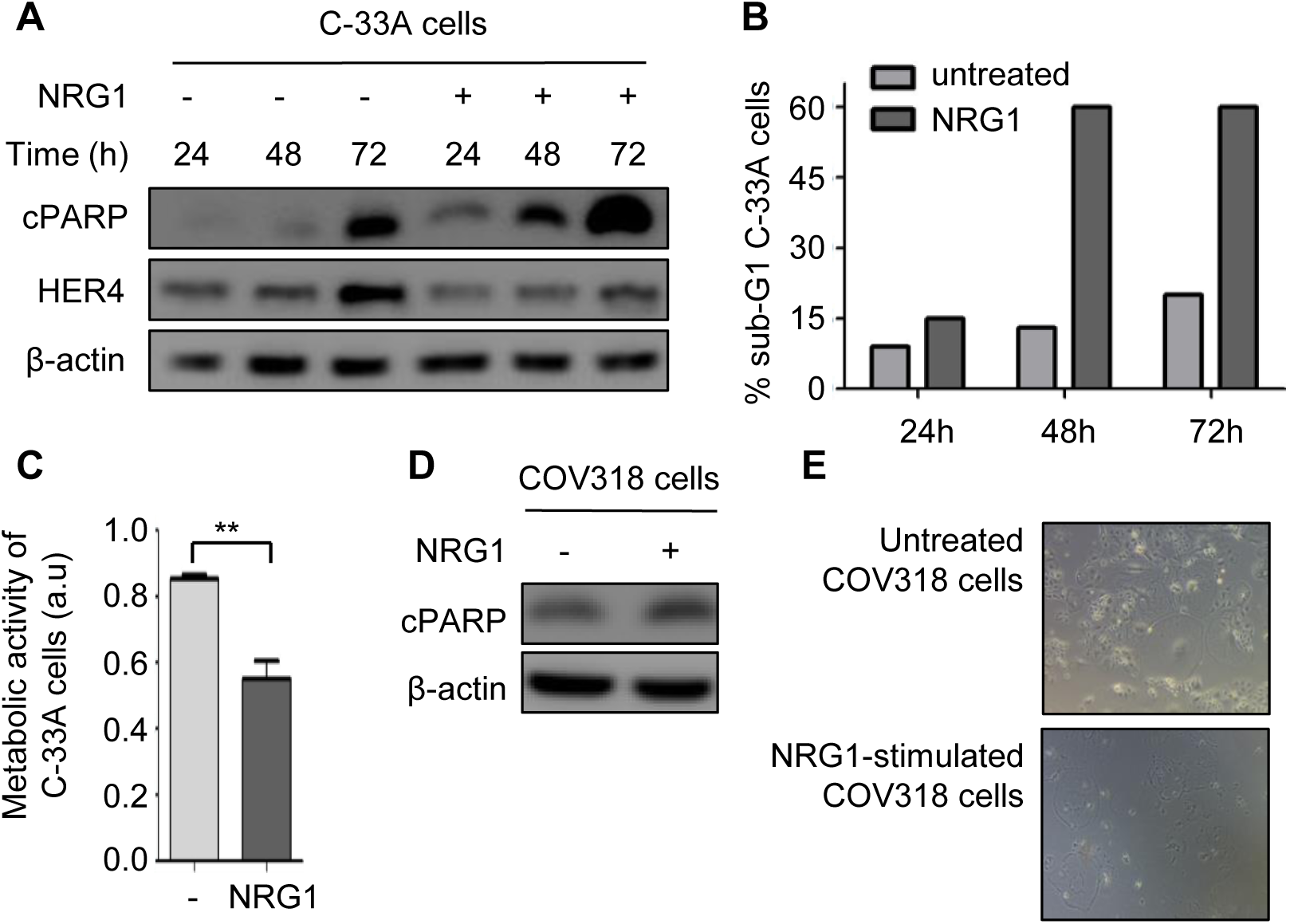
NRG1 induces cell death in HER4-expressing C-33A and COV318 cancer cells. (**A**) C-33A cervical cancer cells were serum-starved for 12h, stimulated with NRG1 (30 ng/ml) for the indicated times and analyzed by western blotting using the appropriate antibodies. (**B**) C-33A cells were treated as described in (A), stained with propidium iodide and analyzed by flow cytometry. (**C**) C-33A cells were serum-starved for 12h and stimulated with NRG1 (30 ng/ml) for 5 days. Metabolic activity was analyzed with the MTS assay. Data are representative of two independent experiments and are presented as mean±SEM. **p<0.01 (unpaired t test). (**D**) COV318 ovary cancer cells were serum-starved for 12h, stimulated with NRG1 (30 ng/ml) for 24h and analyzed by western blotting using the appropriate antibodies. (**E**) Photographs showing the phenotype of unstimulated cells (top) and NRG1-stimulated COV318 cells (bottom).

### 3.2. NRG1 induces cell death through the HER4 JMa/CYT1 isoform

It was previously demonstrated that 4ICD, after cleavage of the JMa/CYT1 or JMa/CYT2 isoform, favors apoptosis [19] To determine which isoform is responsible for cell death after NRG1 stimulation, we transiently-transfected HER3^neg^/HER4^neg^ BT549 TNBC cells with plasmids that encode full-length HER4 JMa/CYT1 or JMa/CYT2. Flow cytometry analysis confirmed HER4 expression and showed that EGFR and HER2 expression were not affected by transfection (Supplementary Fig. S3A). We also checked the expression of both full-length HER4 isoforms by western blotting (Supplementary Fig. S3B), and the absence of expression in mock-transfected BT549 cells. Expression of HER4 JMa/CYT1 was lower than that of JMa/CYT2 (Supplementary Fig. S3B), in accordance with literature data [14, 16]. RT-PCR analysis confirmed that non-transfected BT549 cells did not express any HER4 isoform (Supplementary Fig. S3C).

In basal conditions (medium alone), the expression of each HER4 isoform in BT549 cells was sufficient to induce PARP cleavage at 48h post-transfection. Stimulation with NRG1 sustained PARP cleavage up to 72h in JMa/CYT1-transfected BT549 cells, but not in mock- and JMa/CYT2-transfected cells where PARP cleavage decreased (Fig. 2A). To confirm the opposite roles of HER4 JMa/CYT1 and JMa/CYT2, we overexpressed each isoform in C-33A cells. These cells naturally express the two isoforms at similar levels, but not the JMb isoforms (Supplementary Fig. S3C). NRG1-induced PARP cleavage was comparable in parental and mock-transfected C-33A cells (compare Fig. 2B and Fig. 1A). HER4 JMa/CYT1 overexpression, but not JMa/CYT2 overexpression strongly induced PARP cleavage in unstimulated cells, particularly at 72h (Fig. 2B). Upon NRG1 stimulation, PARP cleavage was amplified from 48h onwards in HER4 JMa/CYT1-overexpressing C-33A cells, whereas it was decreased in JMa/CYT2-overexpressing cells, compared with NRG1-stimulated mock-cells (Fig. 2B).

**Fig. 2.**
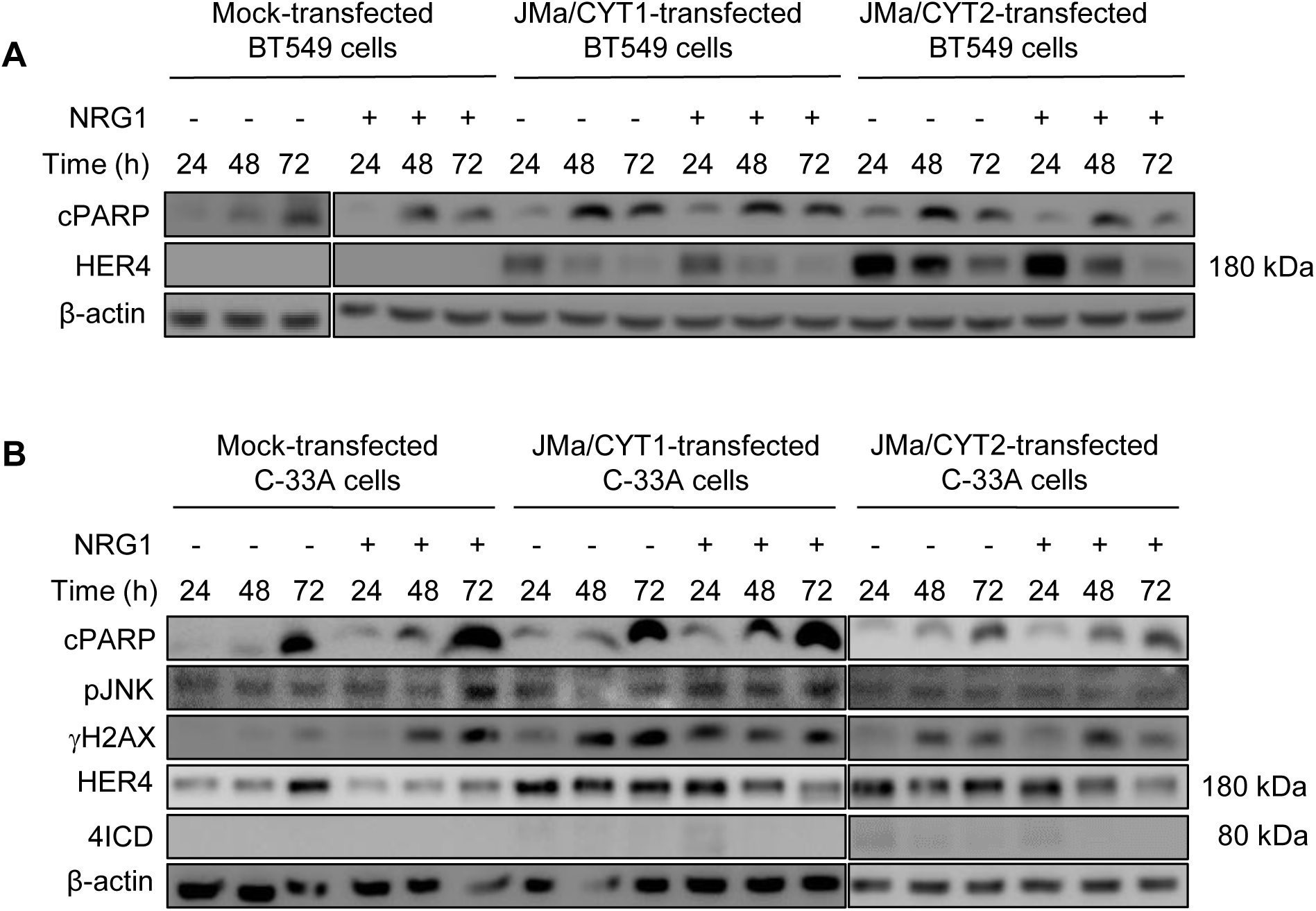
NRG1 induces cell death through the HER4 JMa/CYT1 isoform and JNK/γH2AX signaling. (**A**) BT549 TNBC cells were transfected with plasmids encoding full-length HER4 JMa/CYT1 or JMa/CYT2, or Mock-transfected. At 24h post-transfection, cells were serum-tarved for 12h, stimulated or not with NRG1 (30 ng/ml) for the indicated times and analyzed by western blotting using the appropriate antibodies. (**B**) C-33A cells that naturally express both HER4 JMa isoforms were transfected with plasmids encoding full-length HER4 JMa/CYT1 or JMa/CYT2, or Mock-transfected. At 24h post-transfection, cells were serum-starved for 12h, stimulated or not with NRG1 (30 ng/ml) for the indicated times and analyzed by western blotting using the appropriate antibodies.

### 3.3. NRG1-induced cell death through HER4 JMa/CYT1 occurs via JNK/γH2AX signaling

In HER3^neg^/HER4^pos^ C-33A cells, NRG1 stimulation decreased the cell metabolic activity (Fig. 1C) and promoted DNA fragmentation (Fig. 1B). Therefore, we wanted to identify the NRG1/HER4-related intermediates between mitochondria and PARP. It has been shown that sustained JNK activation induces apoptosis by interacting with mitochondrial proteins [32], and that NRG1 activates JNK through HER4 [33]. Moreover, HER4 is connected with DNA damage [34]. In agreement, NRG1 stimulation of mock-transfected C-33A cells increased JNK phosphorylation and the level of γH2AX (a marker of double-strand DNA breaks) over time (Fig. 2B), compared with unstimulated mock cells. In HER4 JMa/CYT1-overexpressing C-33A cells, JNK phosphorylation and γH2AX expression were induced earlier (24h vs 48h) and persisted until the experiment end (72h) in NRG1-stimulated compared with unstimulated cells (Fig. 2B). Moreover, JNK phosphorylation level was lower in the absence of NRG1. Conversely, in HER4 JMa/CYT2-overexpressing C-33A cells, JNK phosphorylation was not induced in any condition (Fig. 2B), and basal γH2AX expression at 72h was reduced by NRG1 stimulation. These experiments confirmed that HER4 JMa/CYT1 acts as a tumor suppressor upon NRG1 stimulation, whereas JMa/CYT2 acts as a tumor protector.

### 3.4. NRG1 induces 4ICD retention in mitochondria of HER4 JMa/CYT1-transfected cells

As it has been shown that cleaved 4ICD closely interacts with mitochondria [19, 21], we asked whether NRG1-induced cell death could be linked to 4ICD-CYT1 localization in mitochondria. To this aim, we investigated 4ICD localization in mitochondria and cytosol, after subcellular fractionation of JMa/CYT1 and JMa/CYT2-transfected BT549 cells. In non-stimulated cells (medium alone), we detected 4ICD in the mitochondrial fraction of HER4 JMa/CYT1- and JMa/CYT2-transfected BT549 cells (Fig. 3A). We also observed a pool of 4ICD-CYT1, but not of 4ICD-CYT2, in the cytosolic fraction of unstimulated cells. After NRG1 stimulation, full-length HER4 mostly disappeared both in the mitochondrial and cytosol fractions, suggesting increased HER4 degradation or shedding. Similarly, the mitochondrial fraction of 4ICD-CYT2, but not of 4ICD-CYT1 decreased (Fig. 3A). These results suggest that NRG1-induced cell death in HER4 JMa/CYT1-overexpressing cells occurs through 4ICD-CYT1 retention in mitochondria.

**Fig. 3.**
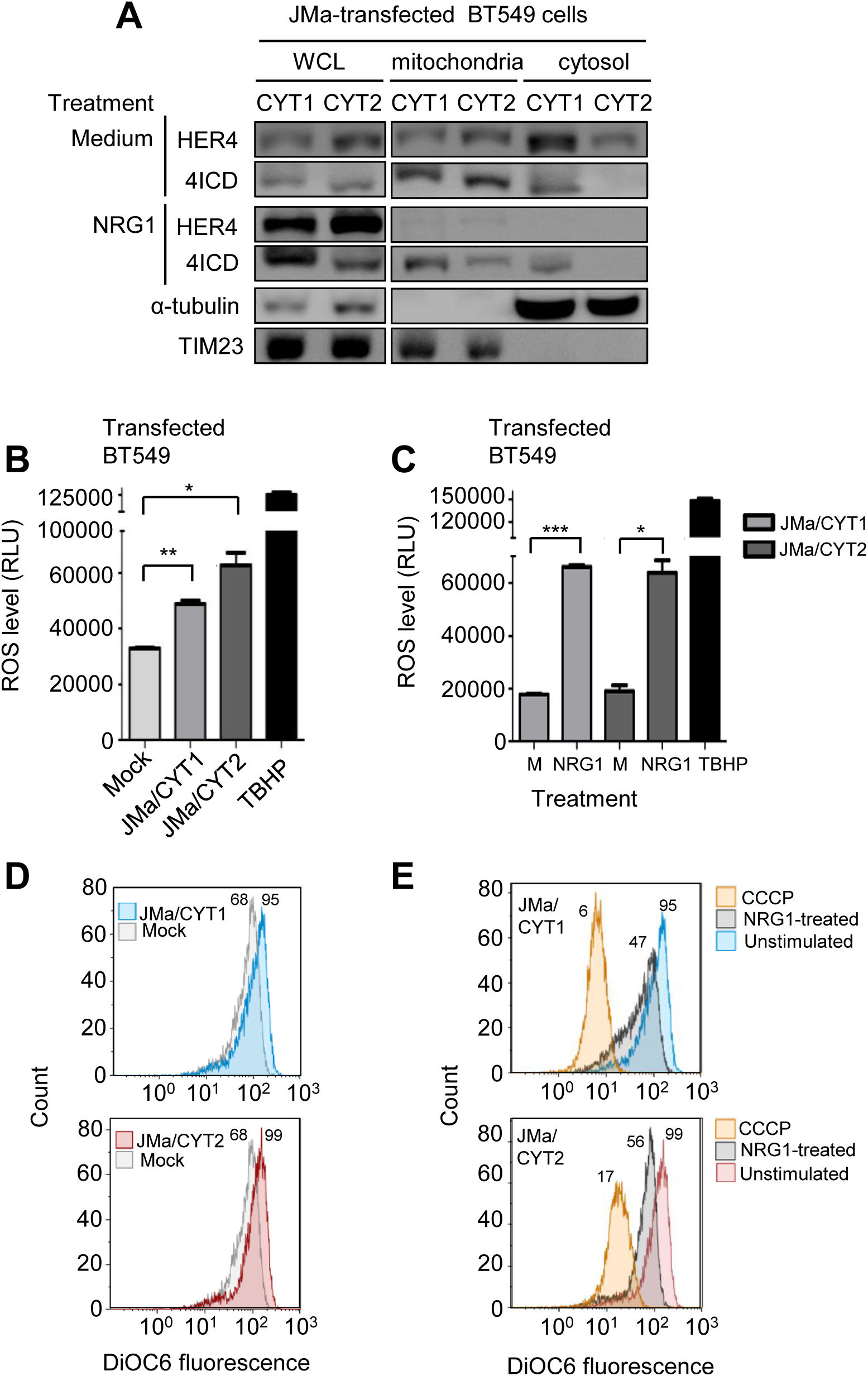
In HER4 JMa/CYT1-transfected cells, NRG1 induces 4ICD retention in mitochondria and ROS production through mitochondrial depolarization. (**A**) Subcellular fractionation of BT549 cells transfected with HER4 JMa/CYT1- or JMa/CYT2-encoding plasmids for 5h, serum-starved for 19h, and stimulated or not (Medium) with NRG1 (30 ng/ml) for 24h. The cytosol and mitochondrial fractions were identified with antibodies against α-tubulin and TIM23, respectively. The localization of full-length HER4 and 4ICD was assessed using the anti-HER4 antibody E200 (Abcam) that recognize full-length HER4 (180 kDa) and 4ICD (80 kDa). WCL: Whole Cell Lysate. (**B**) ROS quantification in HER4 JMa/CYT1- and JMa/CYT2-transfected BT549 cells vs Mock-transfected cells. At 24h post-transfection, cells were serum-starved for 18h, and ROS levels were quantified with the Cellular Reactive Oxygen Species Detection Assay Kit (Abcam). The ROS-inducer TBHP was used as positive control. Data are presented as mean±SEM. *p<0.05, **p<0.01 (unpaired t test). RLU: Relative Luminescence Unit. (**C**) ROS measurement after NRG1 stimulation of HER4 JMa/CYT1- and JMa/CYT2-transfected BT549 cells. At 24h post-transfection, cells were serum-starved for 12h and then stimulated or not (M) with NRG1 (30 ng/ml) for 24h. ROS levels were quantified as in (B). Data are presented as means±SEM. *p<0.05, **p<0.01, ***p<0.001 (unpaired t test). RLU: Relative Luminescence Unit. (**D**) Mitochondrial Membrane Potential (MMP) measurement of HER4 JMa/CYT1 and JMa/CYT2-transfected BT549 cells vs Mock-transfected cells. At 24h post-transfection, cells were serum-starved for 12h and MMP was analyzed by flow cytometry using the DIOC6 probe. (**E**) MMP measurement after NRG1 stimulation of HER4 JMa/CYT1 and JMa/CYT2-transfected BT549 cells. At 24h post-transfection, cells were serum-starved for 12h and then stimulated or not with NRG1 (30 ng/ml) for 24h. MMP was measured as in (D). The mitochondrial oxidative phosphorylation uncoupler CCCP was used as positive control for mitochondrial membrane depolarization.

### 3.5. NRG1 induces ROS production through mitochondrial depolarization

ROS, the main cause of oxidative stress, is mainly produced in mitochondria, particularly upon sustained JNK activation [35]. Oxidative stress caused by ROS overproduction leads to DNA damage and cell death; however, a “sustained and controlled” ROS production could also favor tumor progression [36]. To investigate ROS role in HER4 JMa/CYT1-mediated cell death, we analyzed ROS production in transfected BT549 cells. Without NRG1 stimulation (Fig. 3B), ROS level was significantly higher in HER4 JMa/CYT1- and JMa/CYT2-transfected BT549 cells than in mock-transfected cells, in correlation with 4ICD localization in mitochondria. As positive control, TBHP strongly induced ROS production. NRG1 stimulation significantly increased ROS production in HER4 JMa/CYT1- and JMa/CYT2-transfected BT549 cells, compared with unstimulated cells (only medium) (Fig. 3C).

As ROS is produced in mitochondria through reduction of the mitochondrial membrane potential, we used the DIOC6 probe to monitor the mitochondrial membrane potential. Flow cytometry analysis of DIOC6 labeled unstimulated HER4 JMa/CYT1- and JMa/CYT2-transfected BT549 cells demonstrated an increase of DIOC6 fluorescence signal, indicating mitochondrial membrane hyperpolarization, compared with mock-transfected cells (Fig. 3D). Conversely, NRG1 stimulation reduced DIOC6 labeling, indicating mitochondrial membrane depolarization (Fig. 3E). As positive control of depolarization, CCCP strongly reduced DIOC6 signal intensity. These results suggest that NRG1 induces ROS production through mitochondrial membrane depolarization of HER4 JMa/CYT1-expressing cells and participates in NRG1-induced cell death.

On the basis of these results, we propose a model in which NRG1 stimulation induces cell death through HER4 JMa/CYT1 activation, releasing 4ICD-CYT1, with a role of “cell death inducer”. In mitochondria, 4ICD interacts with effectors, inducing mitochondrial depolarization that leads to ROS overproduction. As NRG1-stimulated HER4 JMa/CYT1 also induces JNK phosphorylation, the JNK pathway could amplify this phenomenon, leading to PARP cleavage, DNA damage, and death of HER4 JMa/CYT1-expressing cancer cells.

### 3.6. Selection and characterization of four anti-HER4 antibodies from the HUSCI phage-display library

Then, to select anti-HER4 antibodies that could mimic HER4 JMa/CYT1 tumor suppression *via* NRG1, we developed a whole cell panning approach in which we used HER4 JMa/CYT1-transfected NIH3T3 cells as targets (after stimulation with NRG1 to pre-activate HER4) and the proprietary phage display scFv library HUSCI (see Methods). We also performed a second phage display screening of the HUSCI library using soluble recombinant HER4 as target. Among the 400 clones screened by flow cytometry and ELISA, we selected and IgG1-reformatted four antibodies: D5, F4, and C6 (selected using the recombinant protein approach), and H2 (selected by whole cell panning). ELISA showed that the four antibodies bound to human HER4 in a dose-dependent manner (Fig. 4A). This binding was confirmed by flow cytometry in C-33A cells (Supplementary Fig. S4A). D5 and C6, but not F4 and H2, cross-reacted with mouse HER4, as shown by ELISA (Supplementary Fig. S4B). The human HER4-specific D5 (10 ng/ml) and C6 (100 ng/ml) antibodies displayed 50% absorbance (EC50) on human HER4, and bound more efficiently than F4 and H2 (EC50 around 1-5 µg/ml) (Fig. 4A). Competition experiments using NRG1 and flow cytometry demonstrated that addition of low NRG1 concentrations (from 0.1 to 1 nM; 3 to 30 ng/ml) did not hinder antibody binding to HER4 (Fig. 4B). D5 and F4 binding to HER4 was partially inhibited by 10 nM NRG1 (60% and 40% inhibition, respectively), but not C6 and H2 binding (Fig. 4B). F4 and H2, but not C6 and D5, still maintained around 40% of residual binding at high NRG1 concentration (100 nM).

**Fig. 4.**
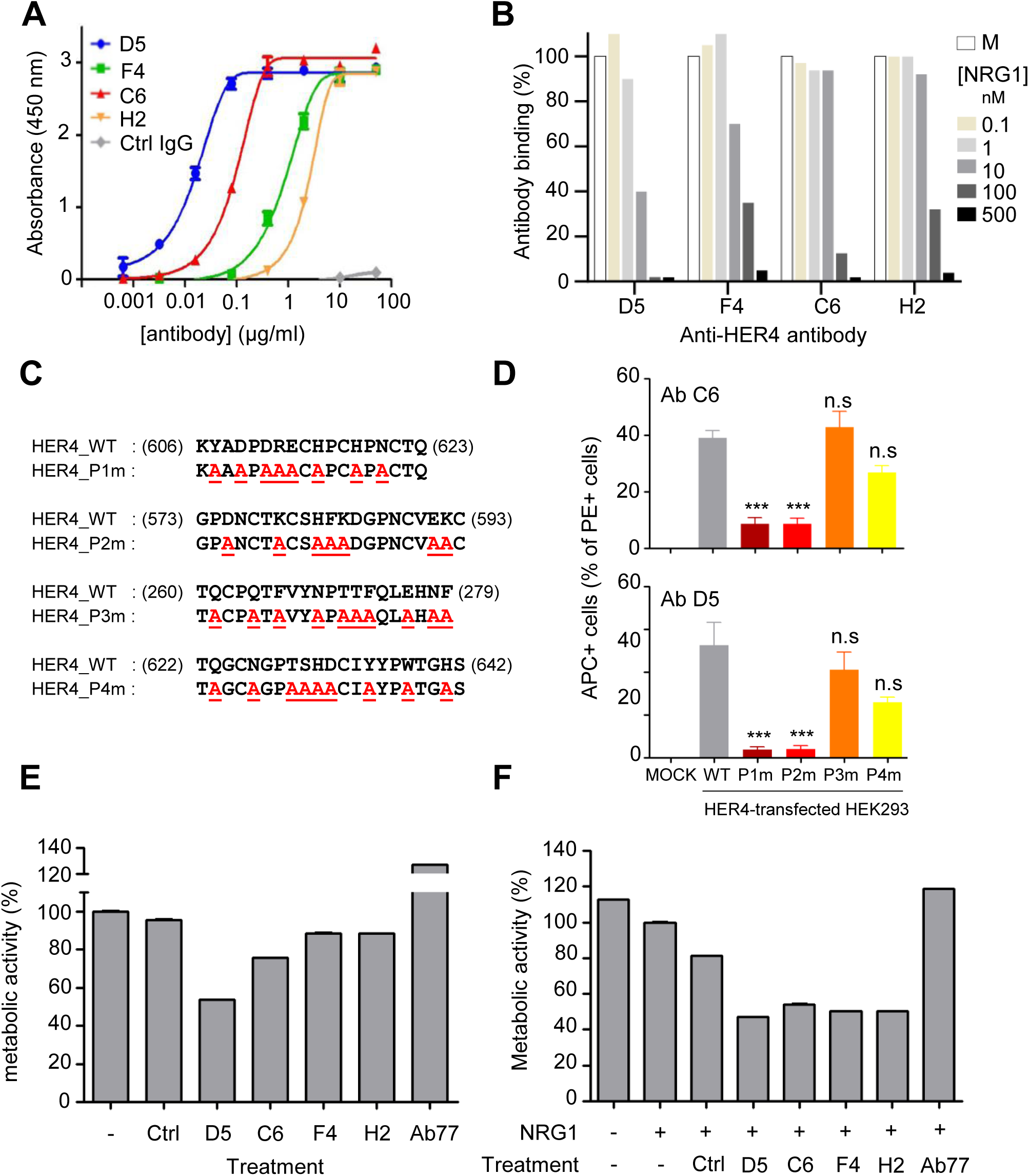
Selection and characterization of four anti-HER4 antibodies from the HUSCI phage-display library. (**A**) ELISA binding of the selected mAbs to human HER4. Wells were coated with 250 ng/ml of recombinant HER4, and 20 μg/ml of mAbs was used as starting concentration (1:10 dilution). Ipilimumab was used as negative control antibody. (**B**) Effect of NRG1 on antibody binding to HER4^pos^ C-33A cells assessed by flow cytometry. Cells were co-incubated with D5, F4, C6 or H2 (15 μg/ml) and various NRG1 concentrations. Results are presented as percentage of binding relative to the binding without NRG1 (M; 100%). (**C**) HER4 variants with mutations within the four predicted areas (HER4_P1 to P4). Only amino-acids from the HER4 protein surface were mutated to alanine, to ensure that HER4 structure was not altered. All wild type (WT) and mutants were N-terminally Flag-tagged and transfected in HEK293 cells, as described in the Materials and Methods section. (**D**) Binding of the anti-HER4 antibodies C6 and D5 in Mock, WT HER4 and HER4 mutant-transfected HEK293 cells. The number of APC+PE+ double-positive cells was quantified by flow cytometry, and normalized to the total number of PE+ cells within the sample. ***p<0.001 (ANOVA test). n.s: non-significant. (**E**) Effect of the four anti-HER4 mAbs on the metabolic activity of C-33A cells. At 24h post-seeding, cells were incubated with D5, F4, C6, H2 or Ab77 (100 μg/ml) for 5 days. Metabolic activity was analyzed using the MTS assay and presented as percentage of the activity in untreated cells (100%). Ctrl: negative control antibody. (**F**) Effect of the four anti-HER4 mAbs on the metabolic activity of NRG1-stimulated C-33A cells. At 24h post-seeding, cells were serum-starved for 12h and then co-incubated with D5, F4, C6, H2 or Ab77 (100 μg/ml) and NRG1 (10 ng/ml) for 5 days. Metabolic activity was analyzed as in (E). Ctrl: negative control antibody. Data in (E) and (F) are representative of two independent experiments and are the mean±SEM.

We predicted the HER4 epitopes of the D5 and C6 antibodies using the MabTope software [31] (Supplementary Fig. S5A-B). We identified four predicted areas that belong to the D5 and C6 putative epitopes: P1 606-623, P2 573-593, P3 260-279 and P4 622-642 (Supplementary Fig. S5B), according to the numbering of the UniProtKB-Q15303 HER4 sequence. We designed HER4 mutants within the four predicted areas (HER4_P1m to P4m) (Fig. 4C) and transfected them in HEK293 cells. We mutated into alanine only surface residues, ensuring that HER4 structure was not altered. Flow cytometry analysis (Supplementary Fig. S5C, and normalized data in Fig. 4D for C6 and D5) showed that the percentage of PE+/APC+ cells, which indicated HER4 expression at the cell surface and antibody binding, was increased in cells transfected with WT HER4 compared with mock-transfected cells. Compared with WT HER4-transfected cells, the percentage of double-positive cells was lower in HER4_P1m- and _P2m-transfected cells, whereas it was not significantly different in HER4_P3m- and _P4m-HER4-transfected cells (Supplementary Fig. S5C and Fig. 4D). These results experimentally demonstrated that the anti-HER4 antibodies D5 and C6 share a common conformational epitope that is located in domain IV, close to the transmembrane region, and restricted to regions 605-620 (part of P1) and 575-592 (part of P2) (Supplementary Fig. S5D).

### 3.7. Effect of the four selected antibodies on the metabolic activity of C-33A cancer cells

To test the efficacy of the four anti-HER4 antibodies, we used an MTS assay to assess the metabolic activity of C-33A cells. Compared with control IgG (Ctrl), the metabolic activity of C-33A cells was reduced by 50% when incubated with D5 and by 15% to 30% in the presence of C6, F4 and H2 (Fig. 4E). Co-incubation with 10 ng/ml of NRG1 increased the C6-, F4- and H2-mediated metabolic activity inhibition to 50% (Fig. 4F), suggesting that NRG1 potentiates the effect of these antibodies. Conversely, NRG1 did not improve D5 inhibitory effect. The anti-HER4 agonist antibody Ab77 increased the metabolic activity of unstimulated and NRG1-stimulated C-33A cells up to 120%, as previously suggested [37].

### 3.8. D5, C6 and H2, but not F4, induce 4ICD cleavage in HER4 JMa/CYT1-transfected cells

To test whether the four anti-HER4 antibodies could potentiate HER4 tumor suppressor activity, we first evaluated their ability to induce 4ICD cleavage in HER4 JMa/CYT1 and JMa/CYT2-transfected BT549 cells. We detected 4ICD by western blotting after incubation of JMa/CYT1-transfected cells with D5, C6 or H2 (Fig. 5A, left panel), suggesting that these antibodies promote JMa/CYT1 4ICD release. We did not observe 4ICD cleavage after incubation of JMa/CYT1-transfected BT549 cells with F4, and also after incubation of JMa/CYT2-transfected cells will the four antibodies (Fig. 5A, right panel). We then asked whether the antibodies activated HER4 through phosphorylation. D5 and C6, but not the anti-HER4 agonist Ab77, strongly induced HER4 phosphorylation on Y1056 in HER4 JMa/CYT1-transfected BT549 cells, a phosphorylation site described as essential for HER4 tumor suppressor function [27] (Fig. 5B). Conversely, Ab77, but not D5 and C6, induced HER4 phosphorylation on Y984 in JMa/CYT1-transfected cells, a site responsible for 4ICD association with STAT5A [38]. No antibody-induced HER4 phosphorylation was observed in HER4 JMa/CYT2-transfected BT549 cells (Fig. 5B). These results suggest that the D5 and C6 mAbs induce HER4 ICD cleavage and HER4 phosphorylation via HER4 JMa/CYT1 activation.

**Fig. 5.**
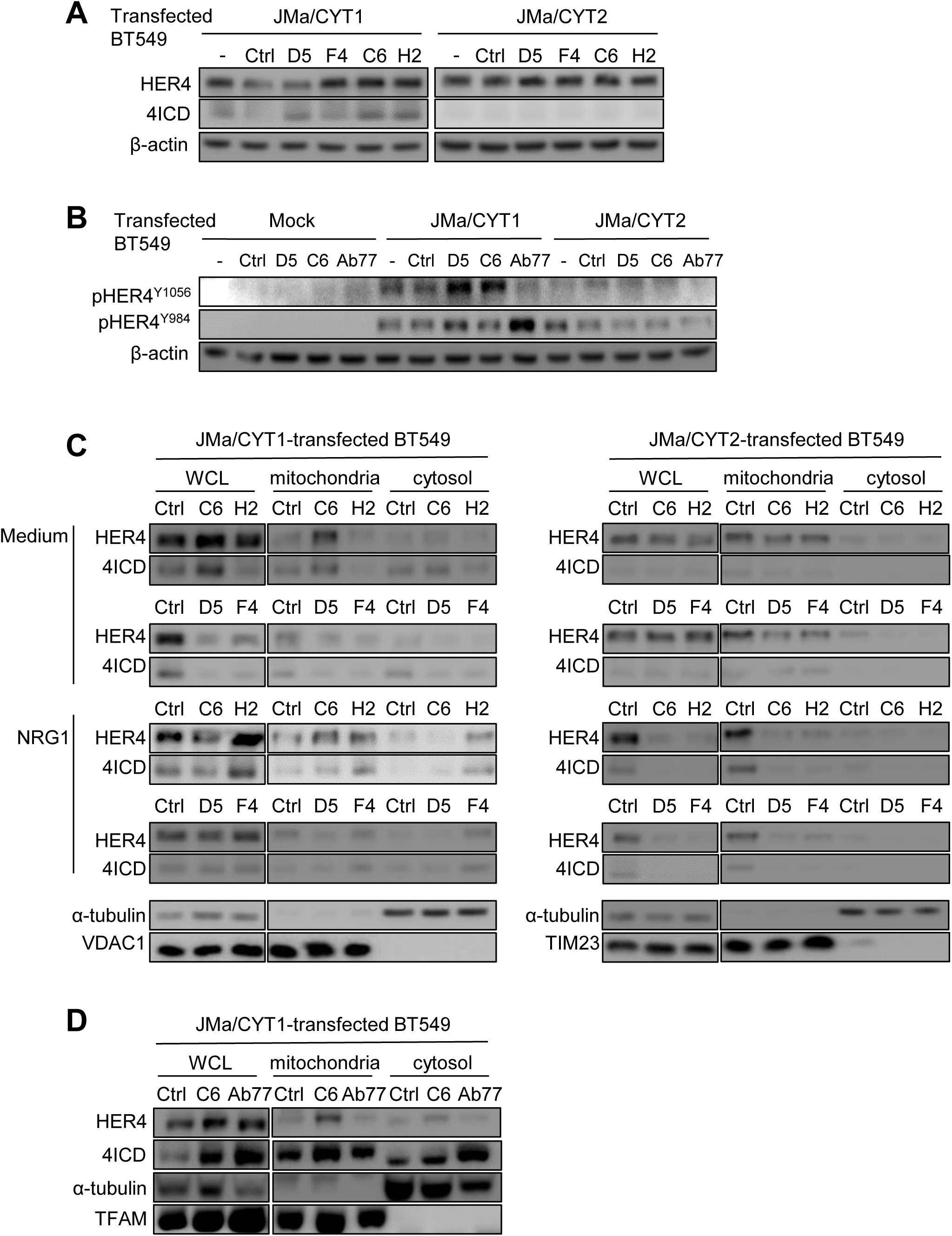
Anti-HER4 antibodies favor 4ICD release and HER4 phosphorylation, leading to 4ICD localization into mitochondria. (**A**) D5, C6 and H2, but not F4 induce 4ICD cleavage from HER4 JMa/CYT1-transfected BT549 cells. At 24h post-transfection with the HER4 JMa/CYT1 or JMa/CYT2 plasmid, cells were incubated with the indicated antibodies (20 μg/ml) for 24h. Full-length HER4 and 4ICD levels were analyzed by western blotting using the anti-HER4 antibody E200 (Abcam) that recognizes full-length HER4 (180 kDa) and 4ICD (80 kDa). (**B**) D5 and C6, but not Ab77, induce HER4 phosphorylation at Y1056 in HER4 JMa/CYT1-transfected BT549 cells. At 24h post-transfection, cells were serum-starved for 12h and incubated with the indicated mAbs (20 μg/ml) for 1h30. HER4 phosphorylation on Y984 and Y1056 was analyzed by western blotting. (**C**) C6 and H2, but not D5 and F4, favor 4ICD retention in mitochondria of HER4 JMa/CYT1-transfected BT549 cells. Cells were transfected with HER4 JMa/CYT1 (left panel) or JMa/CYT2 (right panel) plasmids for 5h, serum-starved for 19h, and then stimulated or not with NRG1 and incubated with 20 μg/ml anti-HER4 antibodies for another 6h. After subcellular fractionation, the cytosolic and mitochondrial fractions were confirmed with antibodies against α-tubulin, and VDAC1 or TIM23, respectively. HER4 and 4ICD localizations were detected using the anti-HER4 antibody E200 (Abcam). WCL: Whole Cell Lysate. (**D**) Subcellular fractionation of HER4 JMa/CYT1-transfected BT549 cells incubated with C6 or Ab77 (20 μg/ml) for 6h. Cells were analyzed as in (B) and (C), with TFAM as mitochondrial fractionation control.

### 3.9. The antibodies C6 and H2, but not D5 and F4, favor 4ICD retention in mitochondria in HER4 JMa/CYT1-transfected cells

We then asked whether the four anti-HER4 antibodies could induce 4ICD retention in mitochondria, as observed following NRG1 stimulation (Fig. 3A). C6 promoted 4ICD retention in mitochondria, as shown by western blotting after subcellular fractionation of HER4 JMa/CYT1-transfected BT549 cells (Fig. 5C; left panels), with and also without co-stimulation by NRG1. H2 promoted 4ICD retention in mitochondria, but only when NRG1 was added (Fig. 5C; left panels). Conversely, C6 and H2 did not have any effect on 4ICD retention in HER4 JMa/CYT2-transfected BT549 cells (Fig. 5C; right panels). This suggests that 4ICD routing to mitochondria is specific to cells that express the HER4 JMa/CYT1 isoform. D5 and F4 did not induce 4ICD retention in mitochondria in HER4 JMa/CYT1- and also JMa/CYT2-transfected BT549 cells (Fig. 5C). We then compared 4ICD localization induced by C6, which inhibited the metabolic activity of C-33A cells, and by the agonist Ab77 antibody, which increased metabolic activity (Fig. 4F). In HER4 JMa/CYT1-transfected BT549 cells, 4ICD-CYT1 retention in mitochondria was higher after incubation with C6 than with Ab77, as expected. Conversely, Ab77 favored 4ICD-CYT1 retention in the cytosol (Fig. 5D). This result shows that these two anti-HER4 mAbs can induce different signaling.

### 3.10. The agonist antibody C6 mimics NRG1-mediated effects on cell death, ROS production, and mitochondrial membrane depolarization

We then asked whether the four HER4-specific antibodies shared properties mediated by NRG1. In C-33A cells, C6 and D5 induced PARP cleavage after 72h of incubation (Fig. 6A) and γH2AX expression, indicating that they favored cell death by promoting double-strand DNA breaks. We confirmed these effects in COV318 cells (Supplementary Fig. S6). C6, but not D5 induced mitochondrial membrane depolarization in HER4 JMa/CYT1-transfected BT549 cells (Fig. 6B), as observed with NRG1 (Fig. 3E). Conversely, incubation of mock- or HER4 JMa/CYT2-transfected BT549 cells with the two anti-HER4 antibodies did not induce any change in mitochondrial membrane potential (Fig. 6B), confirming that depolarization following 4ICD translocation into mitochondria is a specific mechanism to the HER4 JMa/CYT1 isoform. CCCP (positive control) induced mitochondrial membrane depolarization in mock-, JMa/CYT1- and JMa/CYT2-transfected BT549 cells. In addition, compared with mock-transfected cells, ROS production was slightly induced by C6 in HER4 JMa/CYT1-transfected BT549 cells (Fig. 6C, left and middle panels), and by D5 and C6 in JMa/CYT2 cells (Fig. 6C, right panel), suggesting that cancer cell death could be induced also via alternative mechanisms. Incubation with TBHP (positive control) induced ROS production in mock-, JMa/CYT1- and JMa/CYT2-transfected BT549 cells. These results indicate that the anti-HER4 antibody C6 is a cell death inducer that mimics NRG1-mediated effects in cancer cells.

**Fig. 6.**
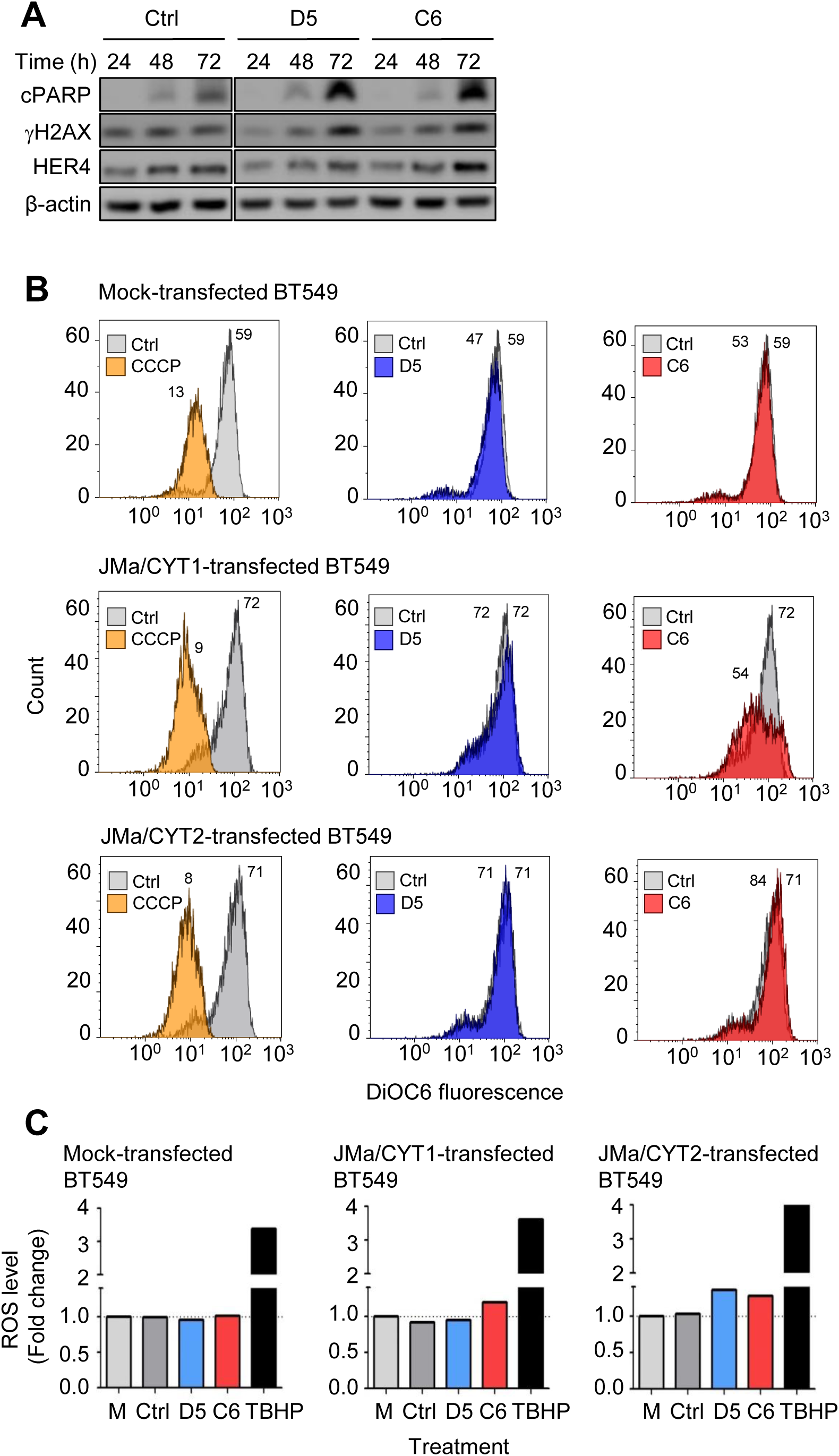
The agonist antibody C6 mimics NRG1-mediated effects on cell death, ROS production, and mitochondrial membrane depolarization. (**A**) C-33A cells were incubated with D5 and C6 (20 μg/ml) for the indicated times and then analyzed by western blotting. (**B**) MMP measurement after incubation with D5 and C6 of Mock-, HER4 JMa/CYT1- and JMa/CYT2-transfected BT549 cells. At 24h post-transfection, cells were serum-starved for 12h and then incubated with antibodies (20 μg/ml) for 24h. MMP was measured using the DIOC6 probe. CCCP was used as positive control. Ctrl: negative control antibody. (**C**) ROS quantification after incubation with D5 and C6 of Mock-, HER4 JMa/CYT1- and JMa/CYT2-transfected BT549 cells. At 24h post-transfection, cells were serum-starved for 12h and then incubated with antibodies (20 μg/ml) for 24h. TBHP was used as positive control. M: medium. Ctrl: negative control antibody.

### 3.11. In vivo the agonist C6 antibody reduces HER4^pos^ ovarian cancer and TNBC cell xenograft growth

Finally, we compared the antitumor efficacy *in vivo* of C6 (NRG1 agonist), D5 and carboplatin (positive control treatment). We xenografted athymic mice with COV434 ovarian cancer cells, C-33A cervical cancer cells and HCC1187 TNBC cells (all HER3^neg^ HER4^pos^). At day 38 post-xenograft (3 days after the end of antibody treatment; day 35), COV434 cell tumor volume was significantly reduced by 33% in D5- and C6-treated mice (p=0.04 and p=0.03, respectively, compared with control IgG), and by 37% in carboplatin-treated animals (p=0.01) (Fig. 7A). At day 41 post-xenograft, the mean tumor volume was reduced by 18% in mice xenografted with C-33A cells and treated with C6 compared with control (IgG), but this difference was not significant (p=0.617) (Fig. 7B). D5 did not affect tumor growth. Conversely, tumor volume was reduced by 59% (at day 38 post-graft) in mice treated with carboplatin (p=0.007). In mice xenografted with HCC1187 cells, tumor volume was reduced by 21.5% (at day 55 post-xenograft, 4 days after the treatment end; day 51) after treatment with C6 compared with control (IgG) (p=0.07) (Fig. 7C). D5 did not have any effect. At day 55 post-graft, tumor size was reduced by 54% in carboplatin-treated mice (p=0.001). These results demonstrate that the agonist anti-HER4 antibody C6 can delay tumor growth in mice xenografted with ovarian cancer or TNBC cells.

**Fig. 7.**
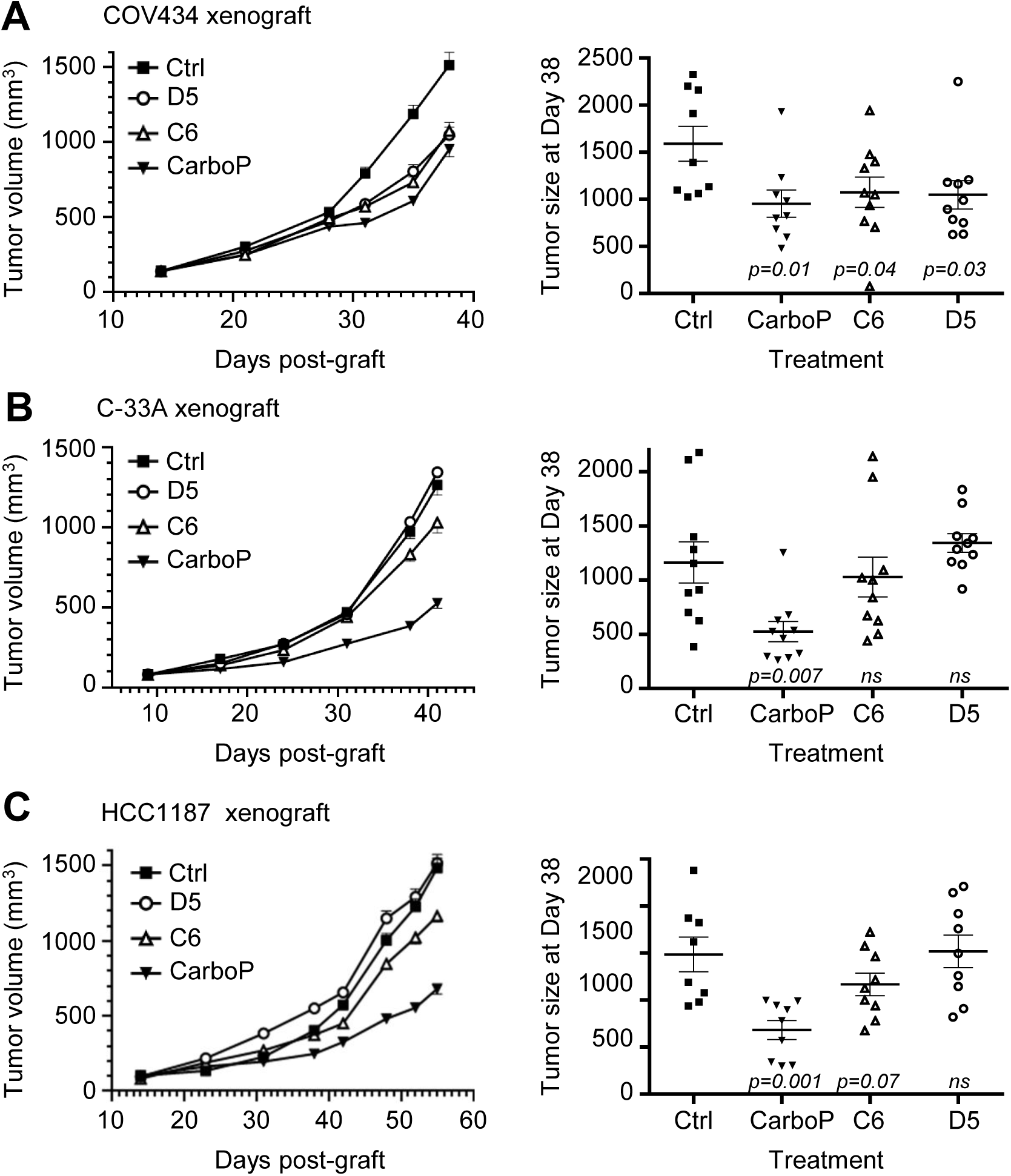
The agonist antibody C6 reduces *in vivo* tumor growth of HER4^pos^ ovarian cancer and TNBC cells. Nude mice (n=10/condition) were xenografted with COV434 ovarian cancer cells (**A**), C-33A cervical cancer cells, or (**B**) HCC1187 TNBC cells (**C**). When tumors reached a volume of 150 mm^3^, mice were treated by intraperitoneal injection of 20 mg/kg of D5 (open white circles) and C6 (open white triangles) (anti-HER4 antibodies), or control antibody (full black squares), twice per week for 4 weeks. Carboplatin (full black triangles; positive control) was used at 60 mg/kg, once per week for 4 weeks. Tumor growth data are presented as the mean tumor volume ± S.E.M. for each group (left panels). The tumor size of each individual mouse is indicated at the end of treatment (right panel). n.s.: non-significant

## 4. Discussion

The development of antagonist mAbs is a classical and effective way to inhibit cancer progression (e.g. anti-EGFR and -HER2 antibodies). The characterization of anti-HER4 mAbs was initially based on the same concept: blocking HER4 activity to kill cancer cells [39]. However, HER4 is unique among the HER family members, with conflicting results concerning the relationship between patient survival and HER4 expression level [40]. Indeed, HER4 displays oncogenic or tumor suppressor activities, depending on the expression levels of its four isoforms [41]. In this context, the previously described anti-HER4 mAbs showed disappointing results because HER4 was targeted as a whole, thus blocking its oncogene and also tumor suppressor activities [42].

In this study, we characterized the pharmacological activity of four new anti-HER4 antibodies by focusing particularly on their effect on each HER4 isoform. The best antibody, C6, is a full agonist molecule that mimics NRG1 effects and promotes cleavage and retention of 4ICD in mitochondria, leading to antibody-induced cell death in HER4 JMa/CYT1-expressing cancer cells. Cell death occurred after ROS production through mitochondrial membrane depolarization, γH2AX expression due to DNA damage, and HER4 phosphorylation at Y1056. *In vivo*, C6 reduced tumor growth in mice xenografted with ovarian cancer or TNBC cells.

In HER3^neg^ HER4^pos^ C-33A cells, NRG1 induces PARP cleavage over time with DNA fragmentation by an unknown mechanism. PARP cleavage after NRG1 stimulation has been described in neurodegenerative diseases, and various cell death mechanisms can potentially be activated through HER4 [43]. We suspect an unconventional cell death mechanism (e.g. necroptosis or parthanatos) through a caspase-independent mechanism that involves endoplasmic reticulum stress. Indeed, protein level was increased after NRG1-induced cell death of HCC1187 cells, suggesting protein synthesis increase, as previously described [44]. Moreover, protein synthesis has been associated with ROS production and JNK activation [45], two events we detected in NGR1-stimulated HER4 JMa/CYT1-expressing cells. Protein synthesis could be also associated with parthanatos, a mitochondrial cell death that implicates ROS, AIF release, and PARP as central mediator [46].

The anti-tumor effect of the C6 antibody is related to HER4 JMa/CYT1 cleavage and formation of a stable active 4ICD fragment located in mitochondria. There are many evidences about HER4 tumor suppressor function through 4ICD, but the exact role of each isoform was never defined. We excluded JMb isoforms because they are not expressed in cancer and they are detected only in some tissues [6]. Using plasmids encoding the two full-length JMa isoforms, we demonstrated that NRG1-induced cell death occurred through HER4 JMa/CYT1. Conversely, HER4 JMa/CYT2 induced cell survival in BT549 cells, a TNBC model that initially does not express HER3 and HER4. Particularly, stimulation by NRG1 increased PARP cleavage in HER4 JMa/CYT1-but not in JMa/CYT2-transfected cells, highlighting the dichotomy between the tumor suppressor JMa/CYT1 isoform and the pro-survival JMa/CYT2 isoform. HER4 JMa/CYT1 acts by activating JNK, relocating 4ICD to the mitochondria and increasing ROS production. Altogether, these events lead to cell death, but we still have to precisely decipher the pathway. We hypothesized that ROS-induced DNA damage could be due to transient mitochondrial membrane potential changes, leading to a cell death mechanism called ROS-Induced ROS Release (RIRR), whereby ROS production is amplified in neighboring mitochondria [47]. Finally, *in vivo*, C6 inhibited growth of HCC1187 tumor cell xenografts (TNBC), possibly through is pro-apoptotic activity. As HER4 is expressed in about 20% of TNBC [41], a pathology still with unmet medical needs, targeting HER4 with agonist antibodies might represent an alternative strategy to treat TNBC.

We used this model as template for HER4 antibody discovery. As NRG1 induces cell death by activating HER4 JMa/CYT1, we tried to potentiate this function without hampering NRG1 action on HER4. To this end, we performed whole cell panning by phage display using NRG1-stimulated HER4 JMa/CYT1-transfected cells to select anti-HER4 mAbs with unique NRG1 agonist/modulator activity. First, NRG1 binding to HER4 differently affects mAb binding to the receptor. Using C-33A cells, we demonstrated that all selected mAbs can bind to HER4 in the presence of 30 ng/ml NRG1, a concentration that induces cancer cell death. Second, the selected mAbs act synergistically with NRG1 to decrease the cell metabolic activity. Third, the epitopes of C6 and D5 are far from the NRG1 binding site [48] on HER4, suggesting that cooperation between HER4-specific mAbs and NRG1 could occur at the cell surface.

Similarly to NRG1, C6 promotes HER4/4ICD localization in mitochondria, mitochondrial depolarization and ROS production, leading to cell death. These HER4 JMa/CYT1-triggered pathways suggest that the 16 aa stretch in CYT1 is crucial for cell death. Interestingly, the previously described anti-HER4 antibody MAb-3 [39] enhances apoptosis in HER4^pos^ non-small-cell lung carcinoma cells, with an increase of sub-diploid cells (a sign of DNA damage). Using the same MAb-3 antibody, Ben-Yosef et al. observed multiple apoptotic cells with pyknotic and fragmented nuclei, karyorrhexis, and loss of cytoplasm in prostate tumor xenograft sections from MAb-3-treated nude mice [49]. Breast cancer frequency is increased in HER4 JMa/CYT1 transgenic mice compared with HER4 JMa/CYT2 transgenic mice, with no apoptosis observed [9]. Thus, cell death induction seems to be a critical point for inhibiting tumorigenesis via HER4 [17]. We demonstrated that the C6 mAb efficiently kills ovarian cancer and TNBC cells *in vitro* by inducing PARP cleavage and DNA damage, and *in vivo* by reducing tumor growth. The anti-tumor effect of this new first-in-class anti-HER4 mAb must be confirmed in additional pre-clinical studies, especially in TNBC.

Finally, as the C6 mAb has a different binding site than NRG1 on HER4, our results suggest that similar pathways can be triggered from different receptor conformations, and indicate that C6 could be an allosteric modulator of HER4. Allosteric modulation is an important mechanism that has been recently adapted from small to large molecules [28, 50]. By acting on the receptor, allosteric molecules can modulate endogenic ligand binding and/or signaling. C6 effect, combined with NRG1 binding on HER4, could exemplify allosteric modulation, with cooperation to initialize new signaling pathways through HER4 that cannot be activated by each molecule on its own. This phenomenon might implicate receptor rearrangement. The finding that C6, D5, Ab77 [37] and mAb1479 [51] have closely related epitopes but very different mechanisms of action suggests that minor differences in conformational changes can induce major differences in signaling and cell fate. This study is the proof-of-concept that biased signaling can be induced with RTK-specific antibodies, as observed with GPCR-targeting molecules. Indeed, the C6 mAb localizes 4ICD in mitochondria, whereas the Ab77 mAb localizes 4ICD in the cytosol of HER4 JMa/CYT1-expressing cells, resulting in different cell fate. Altogether, our observations pave the way to novel mAbs with “biasing properties” for cancer treatment.

## Acknowledgements

We thank S. Bousquié (IRCM) for cell culture and T. David (IRCM) for antibody production. The staff members at the IRCM animal facility, the GenAc and MRI platforms, and the RHEM histology facility are greatly acknowledged. Y. Yarden (Weizmann Institute) is also greatly acknowledged for providing the anti-HER4 antibody Ab77.

## Supplementary data

**Tab. S1.** References of reagents and resources used in this study

**Fig. S1.** Flow cytometry analysis of HER receptor expression in C-33A cervical cancer cells (**A**), and COV318 ovary cancer cells (**B**). Ctrl: negative control antibody.

**Fig. S2.** (**A**) Microphotographs of unstimulated and NRG1-stimulated HCC1187 TNBC cells. (**B**) Adenylate kinase (AK) release from unstimulated and NRG1-stimulated HCC1187 cells. Cells were serum-starved for 12h, and stimulated or not with NRG1 for 72h. Then, cell death was analyzed using an AK-releasing assay. Data are represented as mean ± SEM. **p<0.01 (unpaired t test).

**Fig. S3.** (**A**) Flow cytometry analysis of HER expression in Mock-, HER4 JMa/CYT1- and JMa/CYT2-transfected BT549 cells. Ctrl: negative control antibody. (**B**) Western blot analysis of HER4 and 4ICD expression in parental (wt), mock-, HER4 JMa/CYT1- and JMa/CYT2-transfected BT549 cells using the E200 antibody that recognizes both full-length HER4 (180 kDa) and 4ICD (80 kDa). GAPDH was used as loading control. (**C**) RT-PCR analysis of HER4 isoform expression in non-transfected BT549 (TNBC) cells and C-33A (cervical cancer) cells.

**Fig. S4.** (**A**) Flow cytometry analysis of the binding of the selected antibodies D5, F4, C6 and H2 in C-33A cancer cells. Ctrl: negative control antibody. (**B**) ELISA analysis of the binding of the selected anti-HER4 antibodies to mouse recombinant HER4. Ipilimumab was used as negative control antibody (Ctrl IgG).

**Fig. S5.** D5 and C6 epitope characterization using the MabTope technology. (**A**) Prediction of the best positions for the D5 and C6 epitope using the MabTope *in silico* method. D5 and C6 3D-structures were modeled, docked on HER4 (PDB:2AHX), and 5×10^8^ conformational positions were generated for the complexes. The view of the top 30 ranked predicted conformations for the complex between C6 and D5 (in colors) and the HER4 structure (in grey) is presented. (**B**) Probability for HER4 residue implication in the D5 and C6 epitopes. For each mAb, the frequency of each residue in the epitope was evaluated from the 30 best predicted interfaces. Frequency is represented in blue (darker color indicates higher probability for the residue to be implicated in the epitope). Four regions in HER4 ECD (P1 to P4) appeared to be implicated in mAb binding. (**C**) Flow cytometry analysis of C6 and D5 binding in mock-, WT HER4-, HER4 mutant (P1m-, P2m-, P3m- and P4m)-transfected HEK293 cells. Each HER4 variant harbors a mutation in one of the four predicted regions (HER4_P1m to P4m) (Figure 4C). After cell fixation, membrane HER4 expression was monitored with the PE-coupled anti-Flag antibody (ordinate), and C6 or D5 binding was measured with an APC-coupled anti-IgG antibody (abscissa). An irrelevant IgG was used as control (Ctrl). (**D**) Localization of the regions 575-592 and 605-620 (involved in the conformational epitopes of anti-HER4 antibodies C6 and D5) on the HER4 crystallographic structure (PDB:2AHX).

**Fig. S6.** Western blot analysis of COV318 cells incubated with the D5 and C6 mAbs for the indicated times. Cleaved PARP, γH2AX, HER4 and 4ICD were detected using the appropriate primary antibodies.

