## Supplemental data for "Biasing HER4 Tyrosine Kinase Signaling with Antibodies: Induction of Cell Death by Antibody-Dependent HER4 Intracellular Domain Trafficking"

### Table S1. Reagents and resources

| REAGENT or RESOURCE | SOURCE | IDENTIFIER |
|---|---|---|
| Antibodies | | |
| Anti-HER4 human monoclonal antibody (D5) | This paper | N/A |
| Anti-HER4 human monoclonal antibody (F4) | This paper | N/A |
| Anti-HER4 human monoclonal antibody (C6) | This paper | N/A |
| Anti-HER4 human monoclonal antibody (H2) | This paper | N/A |
| Anti-HER4 mouse monoclonal antibody (Ab77) | Y. Yarden's lab (Chen et al., 1996) | N/A |
| Anti-EGFR monoclonal antibody (cetuximab) | Merck | N/A |
| Anti-HER2 monoclonal antibody (trastuzumab) | Roche | N/A |
| Anti-HER3 monoclonal antibody (clone SGP1) | Santa Cruz Biotechnology | Cat# sc-53279; RRID:AB_1121499 |
| Anti-HER4 monoclonal antibody (clone H4.77.16) | Thermo Fisher Scientific | Cat# MA1-91014; RRID:AB_2099995 |
| Anti-CTLA-4 monoclonal antibody (ipilimumab) (isotype control) | Bristol-Myers Squibb | N/A |
| Mouse irrelevant IgG1 (isotype control) | MP Biomedicals | Cat# 55939 |
| Human irrelevant IgG1 (isotype control) | MP Biomedicals | Cat# 55908 |
| Goat anti-human IgG (Fc specific)-FITC antibody | Sigma-Aldrich | Cat# F9512; RRID:AB_259808 |
| Goat anti-mouse IgG (H+L)- FITC antibody | Millipore | Cat# 12-506; RRID:AB_390186 |
| PE-coupled anti-Flag antibody | Miltenyi Biotec GmbH | Cat# 130-101-576; RRID:AB_2751038 |
| APC-coupled anti-IgG antibody | Miltenyi Biotec GmbH | Cat# 130-093-189; RRID:AB_1036189 |
| Eculizumab | Alexion Pharmaceuticals | N/A |
| Mouse anti-c-Myc antibody HRP-conjugated (clone 9E10) | Santa Cruz Biotechnology | Cat# sc-40; RRID:AB_627268 |
| Mouse anti-c-Myc antibody AlexaFluor647 conjugated | MBL International | Cat# M047-A64; RRID:AB_1953002 |
| Goat anti-human IgG, F(ab')2 fragment specific antibody HRP conjugated | Jackson ImmunoResearch Labs | Cat# 109-036-006 ; RRID:AB_2337590 |
| HER4 (C-term) rabbit monoclonal antibody (clone E200) | Abcam | Cat# ab32375 ; RRID:AB_731579 |
| Anti-cleaved PARP p25 rabbit monoclonal antibody (clone E51) | Abcam | Cat# ab32064 ; RRID:AB_777102 |
| Beta-actin mouse monoclonal antibody (clone 8H10D10) | Cell Signaling Technology | Cat# 3700 ; RRID:AB_2242334 |
| Phospho-SAPK/JNK (Thr183/Tyr185) rabbit monoclonal antibody (clone 81E11) | Cell Signaling Technology | Cat# 4668 ; RRID:AB_823588 |
| Anti-Histone H2A.X, phospho (Ser 139) mouse monoclonal antibody (clone JBW301) | Millipore | Cat# 05-636 ; RRID:AB_309864 |
| Alpha-tubulin mouse monoclonal antibody (clone DM1A) | Cell Signaling Technology | Cat# 3873 ; RRID:AB_1904178 |
| TIM23 mouse monoclonal antibody | BD Biosciences | Cat# 611223 ; RRID:AB_398755 |

#### Table S1. Reagents and resources (continued)

| | | |
|---|---|---|
| Cleaved Caspase-3 (Asp175) rabbit monoclonal antibody (clone 5A1E) | Cell Signaling Technology | Cat# 9664 ; RRID:AB_2070042 |
| Anti-VDAC1 mouse monoclonal antibody | Millipore | Cat# MABN504 ; RRID:AB_2716304 |
| TFAM rabbit monoclonal antibody (clone D5C8) | Cell Signaling Technology | Cat# 8076 ; RRID:AB_10949110 |
| Phospho HER4 (Tyr1056) rabbit polyclonal antibody | Bioss Antibodies | Cat# bs-13094R |
| Phospho HER4 (Tyr984) rabbit polyclonal antibody | Cell Signaling Technology | Cat# 3790 ; RRID:AB_2099879 |
| GAPDH rabbit monoclonal antibody (clone D16H11) | Cell Signaling Technology | Cat# 5174 ; RRID:AB_10622025 |
| Anti-rabbit IgG (whole molecule) goat polyclonal antibody, HRP conjugated | Sigma-Aldrich | Cat# A0545 ; RRID:AB_257896 |
| Goat anti-mouse IgG, F(ab')2 fragment specific antibody HRP conjugated | Jackson ImmunoResearch Labs | Cat# 115-036-072 ; RRID:AB_2338525 |
| **Bacterial and Virus Strains** | | |
| E. coli TG-1 | Agilent | Cat# 200123 |
| E. coli HB2151 | Agilent | Cat# 930684 |
| KM13 helper phage | NEB | Cat# N0315S |
| **Chemicals, Peptides, and Recombinant Proteins** | | |
| Recombinant human NRG1-beta 1/HRG1-beta 1 ECD protein | R&D Systems | Cat# 377-HB-050/CF |
| Recombinant Human HER4 Fc chimera protein | R&D Systems | Cat# 1131-ER-050 |
| Recombinant Mouse HER4 Fc chimera protein | R&D Systems | Cat# 4387-ER-050 |
| Carboplatin | Merck | N/A |
| DiOC6 intracellular probe | Abcam | Cat# ab189808 |
| Propidium iodide | Sigma-Aldrich | Cat# P4170 |
| RNase A | Qiagen | Cat# 19101 |
| CCCP Mitochondrial Oxidative Phosphorylation Uncoupler | Abcam | Cat# ab141229 |
| Trypsin, TPCK treated | Thermo Fisher Scientific | Cat# 20233 |
| Polyethylenimine Linear, MW 25000 (PEI) | Polysciences | Cat# 23966-1 |
| Metafectene | Biontex Laboratories | Cat# T020 |
| **Critical Commercial Assays** | | |
| CellTiter 96 Aqueous One Solution Cell Proliferation Assay (MTS) | Promega | Cat# G3581 |
| ToxiLight Non-Destructive Cytotoxicity Bioassay Kit | Lonza | Cat# LT07-217 |
| DCFDA/H2DCFDA Cellular Reactive Oxygen Species Detection Assay Kit | Abcam | Cat# ab113851 |
| Annexin A5-FITC/7-AAD Kit | Beckman Coulter | Cat# IM3614 |
| Cytofix/Cytoperm Fixation/Permeabilization Solution Kit | BD Biosciences | Cat# 554714 |
| **Experimental Models: Cell Lines** | | |
| Human: C-33A | ATCC | Cat# CRM-HTB-31 ; RRID:CVCL_1094 |
| Human: COV318 | ECACC | Cat# 07071903 ; RRID:CVCL_2419 |
| Human: HCC1187 | ATCC | Cat# CRL-2322 ; RRID:CVCL_1247 |
| Human: BT549 | ATCC | Cat# HTB-122 ; RRID:CVCL_1092 |
| Human: COV434 | ECACC | Cat# 07071909 ; RRID:CVCL_2010 |
| Human: HEK293T | ATCC | Cat# CRL-3216 ; RRID:CVCL_0063 |

Table S1. Reagents and resources (continued)

| | | |
|---|---|---|
| Mouse: NIH3T3 | ATCC | Cat# CRL-1658 ; RRID:CVCL_0594 |
| Experimental Models: Organisms/Strains | | |
| Mouse: Hsd athymic nude mice | Envigo | N/A |
| Oligonucleotides | | |
| JMa/CYT1 amplification: forward *GGCATTCCACTTTACCACA* reverse *GGGCTGTGTCCAATTTCACT* | This paper | N/A |
| JMa/CYT2 amplification: forward *GGCATTCCACTTTACCACA* reverse *CGGTATACAAACTGGTTCCTATTCGAGTCAATTC* | This paper | N/A |
| JMb/CYT1 amplification: forward *CAAGTATTGAAGACTGCATCGG* reverse *GGGCTGTGTCCAATTTCACT* | This paper | N/A |
| JMb/CYT2 amplification: forward *CAAGTATTGAAGACTGCATCGG* reverse *CGGTATACAAACTGGTTCCTATTCGAGTCAATTC* | This papier | N/A |
| Recombinant DNA | | |
| pCMV_HER4 | This paper | N/A |
| pEZY3 vector | (Guo et al., 2008) | Addgene plasmid: #18672 |
| pEZY3_JMa/CYT1 | This paper | N/A |
| pEZY3_JMa/CYT2 | This paper | N/A |
| HER4_WT | This paper | N/A |
| HER4_P1m | This paper | N/A |
| HER4_P2m | This paper | N/A |
| HER4_P3m | This paper | N/A |
| HER4_P4m | This paper | N/A |
| Software and Algorithms | | |
| MabTope | (Bourquard et al., 2018) | www.mabsilico.com |
| Prism Software v5.1 | GraphPad Software | www.graphpad.com |
| Kaluza Analysis software | Beckman Coulter | www.beckman.com |
| ImageJ | (Schneider et al., 2012) | www.imagej.nih.gov |
| FlowJo software | FlowJo, LLC | www.flowjo.com |
| Other | | |
| T-18 digital ULTRA TURRAX | IKA | Cat# 0003720000 |
| Nalgene Centrifuge Tubes | Thermo Fisher Scientific | Cat# 3138-0030 |

**A** C-33A cervical cancer cells

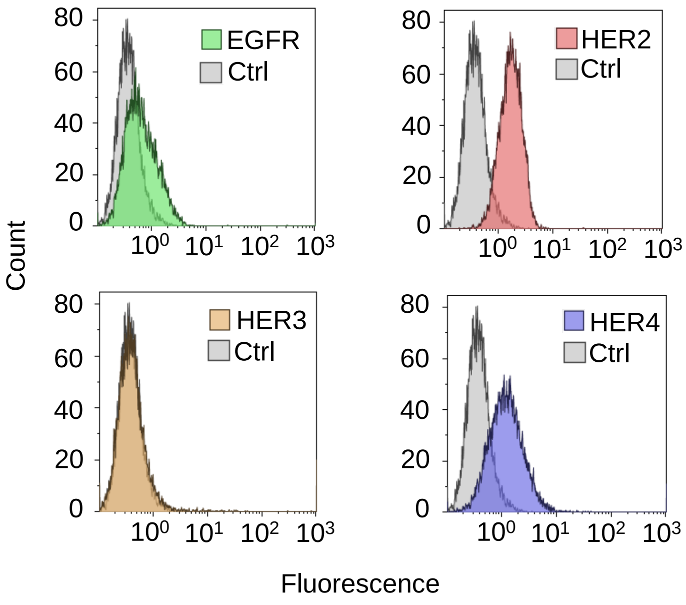


**B** COV318 ovary cancer cells

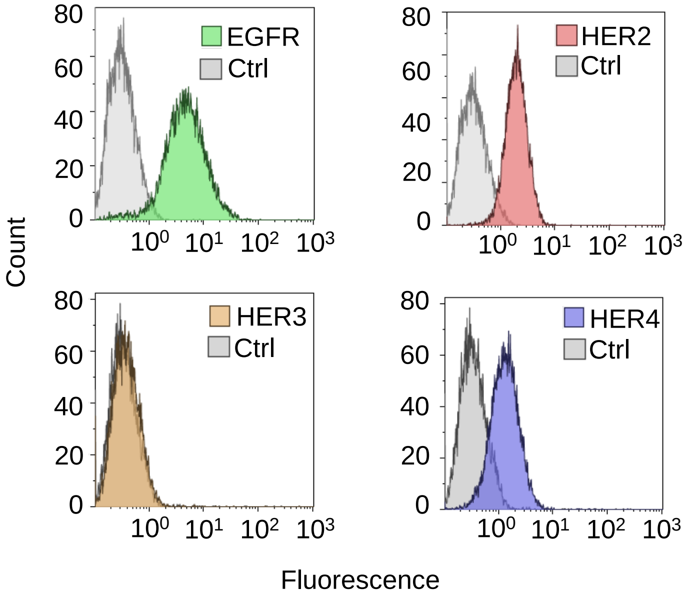


Figure S1. Receptor expression of C-33A and COV318 cancer cells

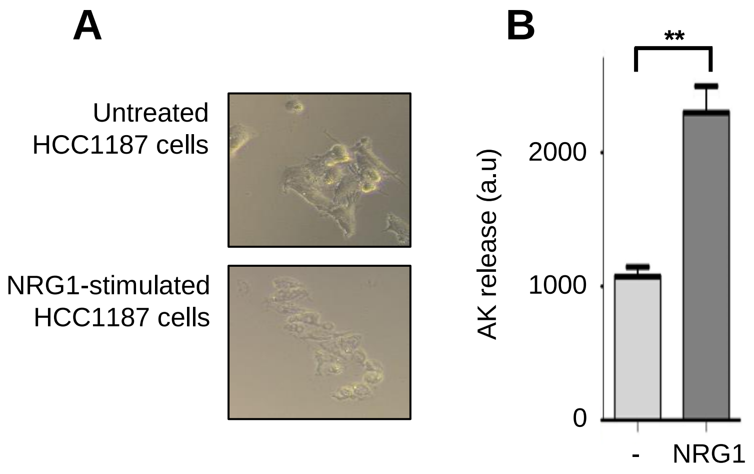


Figure S2. NRG1 induces a « broken » phenotype of HCC1187 cells (A), and reduces Adenylate cyclase release in the supernatant of HCC1187 cells (B). **p<0.01

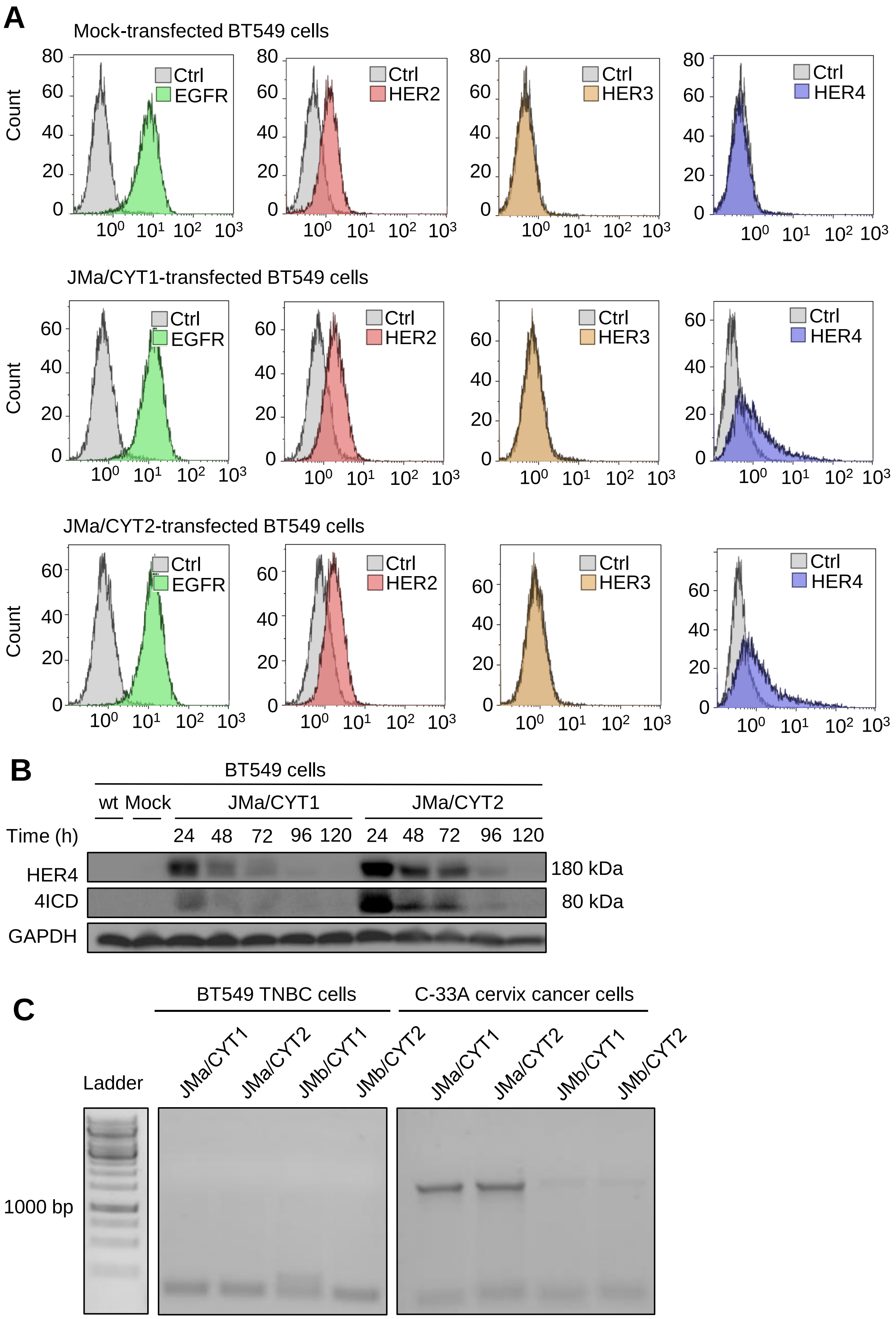


Figure S3. (A) Flow cytometry receptor expression of transfected BT549 cancer cells. (B) Western blot expression of HER4 and 4ICD in wild-type and transfected BT549 cells. (C) RT-PCR of HER4 isoforms in C-33A and BT549 wild-type cancer cells.

A

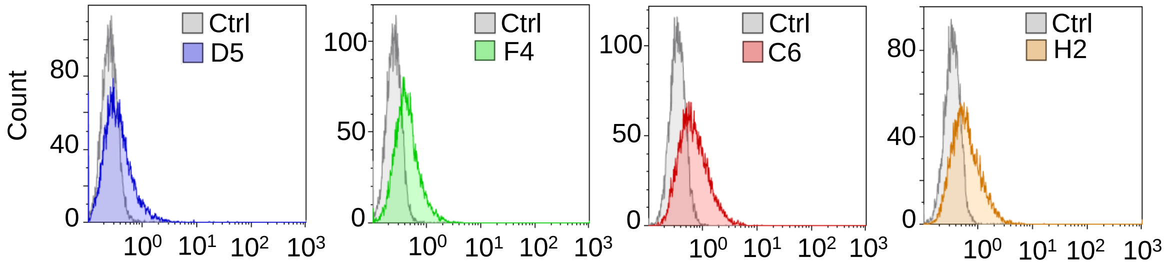


B

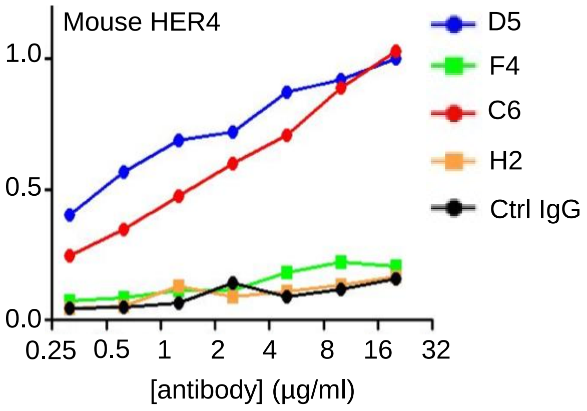


Figure S4. Flow cytometry binding of selected antibodies to membrane HER4 on C-33A cells (A). ELISA antibody binding to mouse HER4 (B).

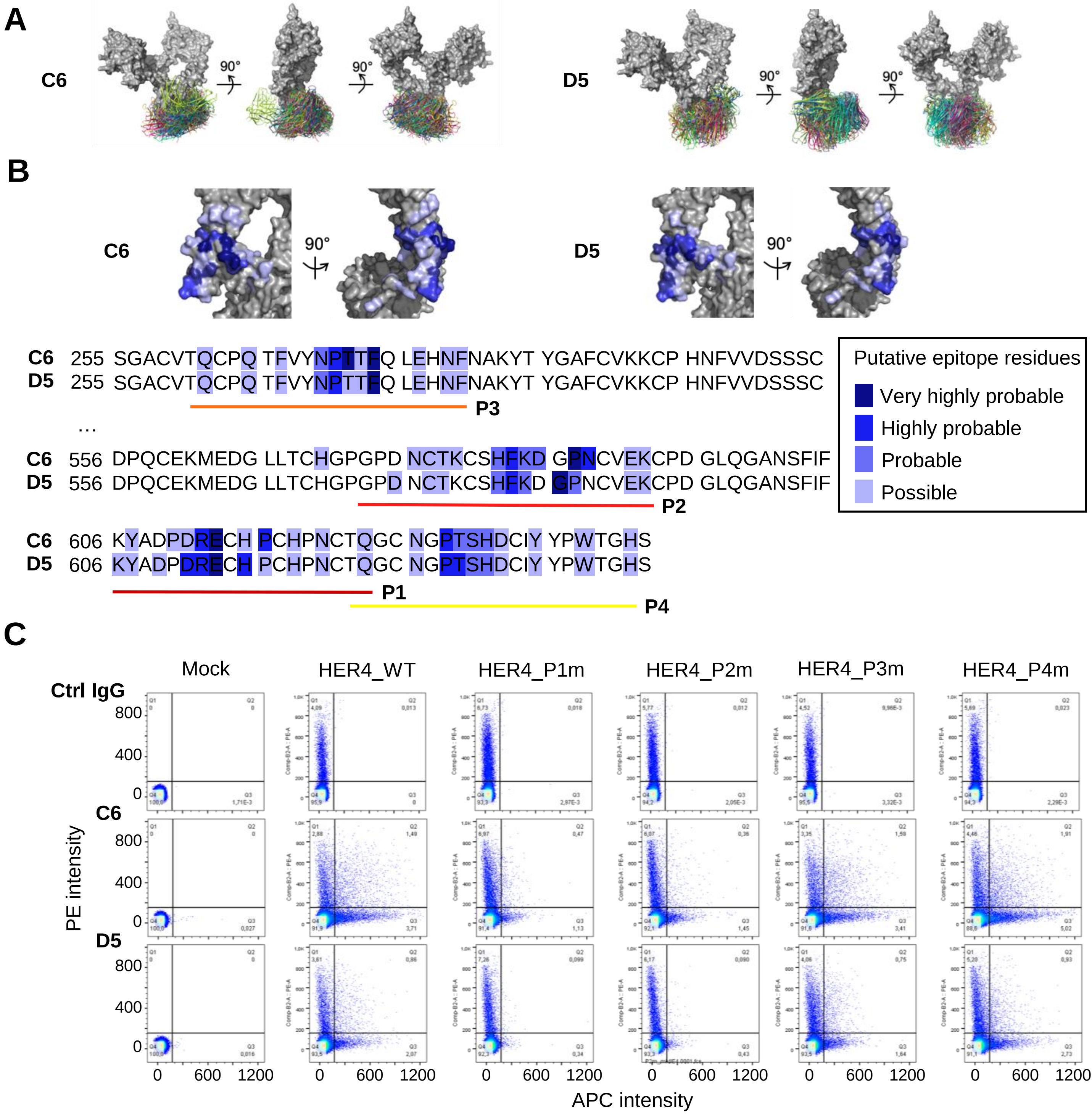


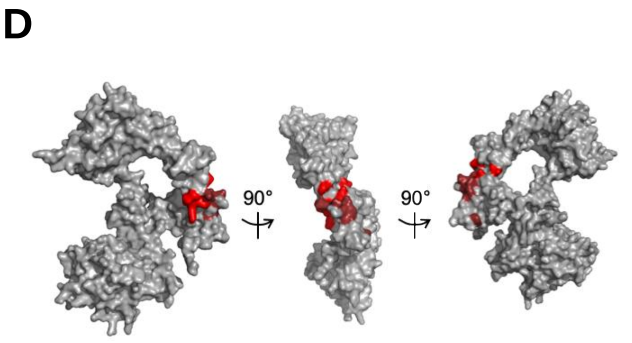


Figure S5. Epitope characterization of C6 and D5 antibodies using the MabTope technology. (A) View of the top 30 ranked predicted conformations for the complex between antibodies C6 and D5 and the HER4 receptor. (B) Residues predicted to belong to the epitopes and selected sequences P1, P2, P3 and P3 for flow cytometry validation. (C) Flow cytometry binding of C6 and D5 antibodies on mock-, wild-type HER4-, mutated HER4 P1m-, P2m, P3m and P4m-transfected HEK293 cells. After cell fixation, membrane HER4 expression was monitored in ordinates with a PE-coupled anti-Flag antibody, and C6 or D5 binding was measured in the abscissa with an APC-coupled anti-IgG antibody. An irrelevant IgG was used as control (Ctrl). (D) Amino-acids corresponding to the shared epitope of C6 and D5 are reported on HER4 structure.

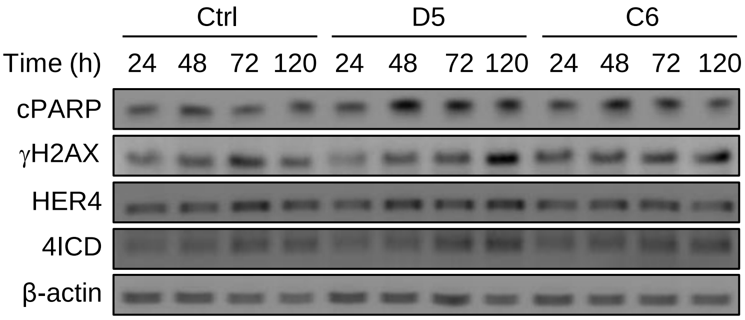


Figure S6. Anti-HER4 antibodies induce PARP cleavage and γH2AX increase in COV318 cells.
